# Stellar quality control for single-cell image-based profiling with coSMicQC

**DOI:** 10.1101/2025.10.14.682427

**Authors:** Jenna Tomkinson, Dave Bunten, Gregory P. Way

**Affiliations:** Department of Biomedical Informatics, University of Colorado Anschutz, Aurora, CO 80045, USA

## Abstract

Over the past twenty years, high-content imaging has transformed our ability to measure cell phenotypes. The need to bioinformatically process these phenotypes led to the development of a research field called image-based profiling. However, because the standard image-based profiling approach involves averaging data, single-cell quality control (QC) has been historically ignored. The conventional approach of aggregating single cells into bulk profiles conserves computational resources and reduces, but does not completely remove, the impact of low-quality single cells. As software scalability improves, researchers are increasingly turning to single-cell image-based profiling to reveal important signals of phenotypic heterogeneity. Therefore, this evolution toward single cells compels single-cell QC standards to ensure that observed morphology differences are driven by biology and not technical interference. We address these challenges with coSMicQC (Single cell Morphology Quality Control), a reproducible Python package with comprehensive tutorials that supports systematic, human-in-the-loop filtering of low-quality single cells. CoSMicQC integrates seamlessly into standard image-based profiling protocols, providing an interactive, Jupyter-compatible user interface to set thresholds and flag technical outliers. Applied to four real-world datasets, coSMicQC achieves data-quality gains comparable to labor-intensive manual annotation, at a fraction of the effort. We show how coSMicQC optimizes assay conditions, outperforms the alternative outlier detector PyOD at single-cell phenotype classification, detects mycoplasma contamination, and rescues lead compounds in a large-scale drug screen that would otherwise have been missed. Overall, coSMicQC is a reliable, scalable method for removing technical outliers, reducing noise, and strengthening image-based profiling insights.

## Introduction

Since the first high-content imaging (HCI) framework was developed over two decades ago, image analysis and image-based profiling have become essential tools for scientific discovery.^1^ Cell Painting is among the most widely used HCI assays, enabling rich measurement of cell phenotypes.^2,3^ Current methodologies of extracting cell morphology features from HCI are Fiji^4,5^, SPACe^6^, Napari^7^, QuPath^8^, and most commonly, CellProfiler.^9^ These methods extract interpretable measurements (e.g., area, shape, texture, intensity, etc.) that require preprocessing before hypothesis testing or other analysis.^10,11^ Open-source software for preprocessing the image analysis outputs includes CytoTable^12^, Pycytominer^13^, BioProfiling.jl^14^, and scMorph^15^. Most image-based profiling pipelines skip single-cell QC, because (1) the canonical averaging procedures reduce the influence of individual cells^16–19^ and (2) QC might also remove single cells with interesting phenotypes and not just technical outliers.^10^ QC is a well-established step in single-cell RNA-seq workflows because it improves data reliability by removing cells with low-quality reads.^20–22^ Despite robust examples of the importance of single-cell QC in RNA-seq, single-cell QC for image-based profiling is not standard practice. Errors derived from image acquisition persist across all image-based profiling experiments, as segmenting microscopy images remains an ongoing challenge.^23^

Recent advances in software and computational processing capabilities have made single-cell microscopy analyses increasingly common, enabling researchers to investigate cell heterogeneity and morphology differences among subpopulations.^11,24–28^ Moreover, with increased single-cell analyses, recent software has developed single-cell QC functionality for microscopy data. For example, CyLinter^29^, which was developed for multiplexed tissue imaging, removes technical artifacts (defined as regions of tissue damage, small antibody aggregates, illumination artifacts, etc.) from single-cell spatial profiles. CyLinter uses predefined metrics (e.g., nuclei intensities and areas) alongside Napari^30^ as a human-in-the-loop interface, enabling users to identify and filter regions of interest (ROIs) containing technical artifacts. However, CyLinter was designed specifically for multiplex tissue imaging, requiring domain-specific input files. For example, CyLinter expects tabular metadata detailing which molecular markers are activated across imaging cycles, which do not apply to the non-spatial, high-content imaging assays used in image-based profiling.

Some QC methodologies have been developed for HCI readouts, including SPACe^6^, which performs both feature extraction and processing, and includes a module for single-cell QC. SPACe uses negative control cells as a reference and compares feature distributions between treated cells to identify “abnormal” single-cell phenotypes. Based on these abnormal phenotypes (or simply low cell counts), SPACe drops whole wells from analysis. This approach does not differentiate technical artifacts from biological phenotypes, but rather considers all extreme differences within the feature space as artifacts. Similarly, scMorph^15^, which is software to process single-cell morphology profiles, utilizes PyOD^31,32^ to detect single-cell outliers. scMorph implements PyOD’s unsupervised empirical cumulative distribution-based outlier detection (ECOD) algorithm with cumulative distribution functions to determine which cells are outliers. Rezvani et al. also uses PyOD, specifically histogram-based outlier scoring, to remove cells that violate normal distributions.^33^ They found that applying cell-level outlier detection outperformed linear regression in improving profile quality. Other modern methodologies of performing cell-level QC include training supervised classifiers to predict which cells are technical artifacts.^34^ On the opposite side of the same coin, Shpigler et al. uses outlier single-cells to identify anomalous cell phenotypes.^35^ This method was designed primarily for representation learning and not QC. A disadvantage of all these methods is that they cannot distinguish whether a cell is an outlier due to an interesting phenotype or a technical issue. The human-in-the-loop approach we take in coSMicQC allows us to filter for technical issues only, a quality we demonstrate in the applications below. Critically, with the exception of scMorph, all of these methods lack installable software that integrates seamlessly into existing high-content microscopy pipelines.

To remove technical single-cell outliers while retaining cells with interesting phenotypes (even within the same well), we developed a reproducible Python package called coSMicQC.^36^ The software is seamlessly compatible with the existing image-based profiling infrastructure developed in the Cytomining community. As input, coSMicQC expects unprocessed, single-cell morphology features harmonized by CytoTable.^12^ CytoTable processes outputs from multiple feature extraction software including CellProfiler, DeepProfiler^37^, and IN Carta. As a result, coSMicQC is not limited to one methodology of feature extraction. The only caveat is that coSMicQC expects the user to have prior knowledge of what each feature represents in order to define interpretable filtering conditions. CoSMicQC uses these features to flag (or filter) poor-quality single-cell segmentations that commonly arise from technical issues such as over-or under-segmentation. Here, we apply coSMicQC to four datasets to demonstrate the impact of single-cell QC. coSMicQC optimizes assay conditions and performs comparably to manual annotation. It also significantly improves machine learning model performance in predicting whether a cardiac fibroblast cell came from a patient with heart failure, matching PyOD ECOD’s performance while filtering far fewer genuine biological cells. We also show that coSMicQC detects mycoplasma contamination. When integrated into traditional image-based profiling pipelines that aggregate single cells into well profiles, coSMicQC has little impact on supervised mechanism-of-action (MOA) prediction, but it rescues 50 lead compounds across all doses, substantially altering rankings in a high-throughput drug screen that uses aggregated bulk profiles. In summary, coSMicQC offers a simple, fast, and seamless methodology within a reproducible software package for detecting poor-quality single cells for image-based profiling.

## Results

### Integrating coSMicQC into standard image-based profiling workflows

We define “outlier” as a technical abnormality caused by improper segmentation. For coSMicQC, we do not consider an outlier as one driven by an interesting and genuine single-cell phenotype. We find that the most common outliers include: (1) segmenting single cells or nuclei into multiple objects (“over-segmentation”), (2) segmenting multiple cells or nuclei together as one object (“under-segmentation”), and (3) segmenting background or debris (“mis-segmentation”).^38^ We developed coSMicQC to flag and/or filter these outliers, thereby improving the accuracy and interpretability of image-based profiling results. CoSMicQC fits into the traditional Cytomining image-based profiling workflow (https://github.com/cytomining) right after CytoTable harmonizes single-cell profiles derived from various image analysis tools like CellProfiler, but before Pycytominer processing (**Figure 1**). CoSMicQC addresses a gap in the high-content screening field where existing single-cell QC approaches do not explicitly distinguish between technical and biological sources of outliers. CoSMicQC differs from other approaches by focusing on selecting a subset of morphology features to find segmentation-derived technical outliers while reducing the removal of interesting phenotypes (**Table 1**).

**Figure 1.**
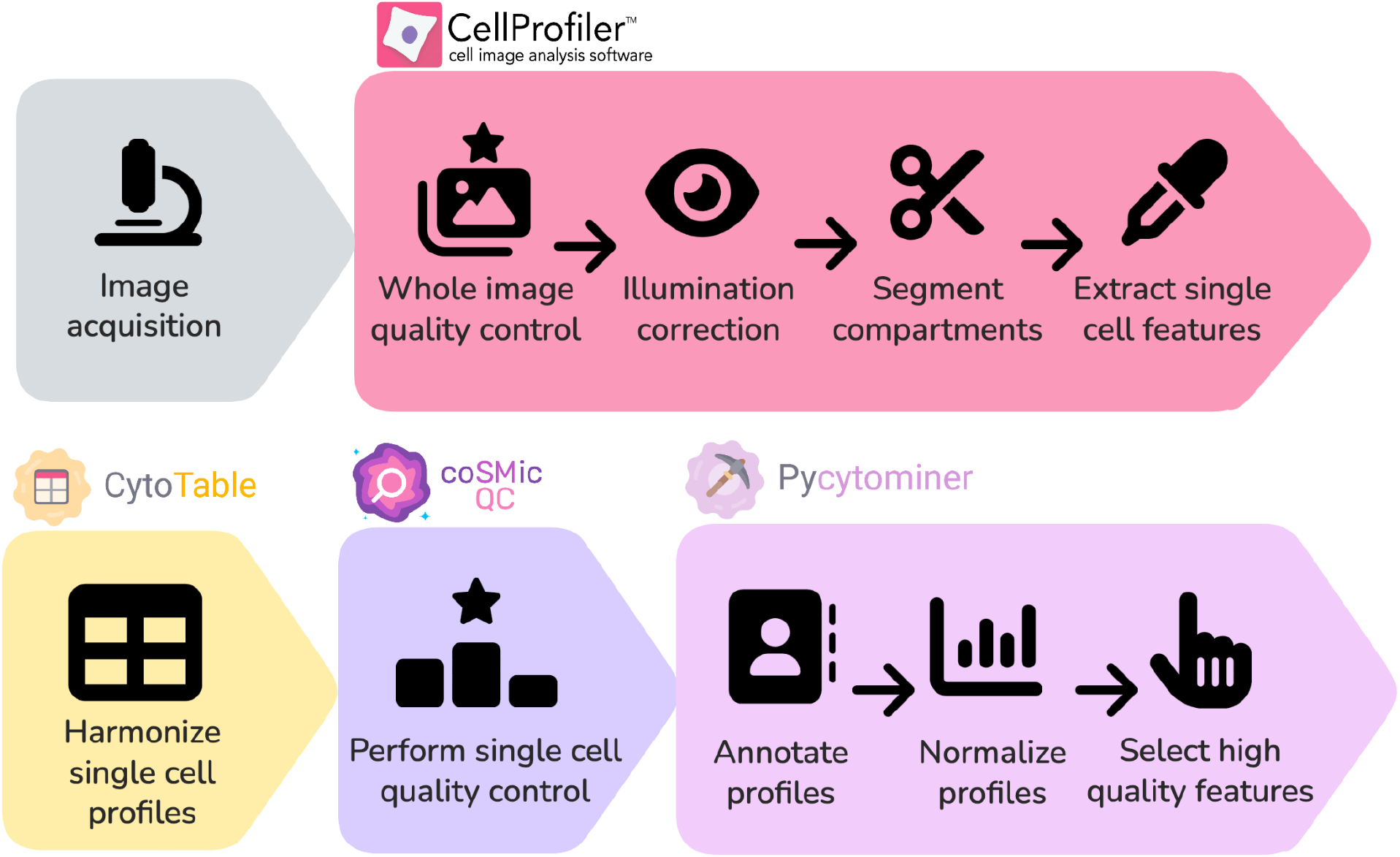
coSMicQC fits seamlessly within the open-source Cytomining image-based profiling workflow. Image-based profiling starts after microscopy image acquisition. The traditional pipeline starts with CellProfiler (or other feature extraction software compatible with CytoTable), which processes raw images through multiple steps, including whole-image QC, illumination correction, segmentation, and morphology feature extraction. Next, CytoTable harmonizes CellProfiler SQLite or CSV output data, which represent the single-cell morphology features extracted per segmented compartment. CytoTable then harmonizes these data into Parquet or AnnData formats as harmonized single-cell profiles. CoSMicQC occurs between CytoTable harmonization and Pycytominer processing. CoSMicQC flags and/or filters low-quality single-cells. Users export the data and continue processing high-quality single cells with Pycytominer.

**Table 1:**
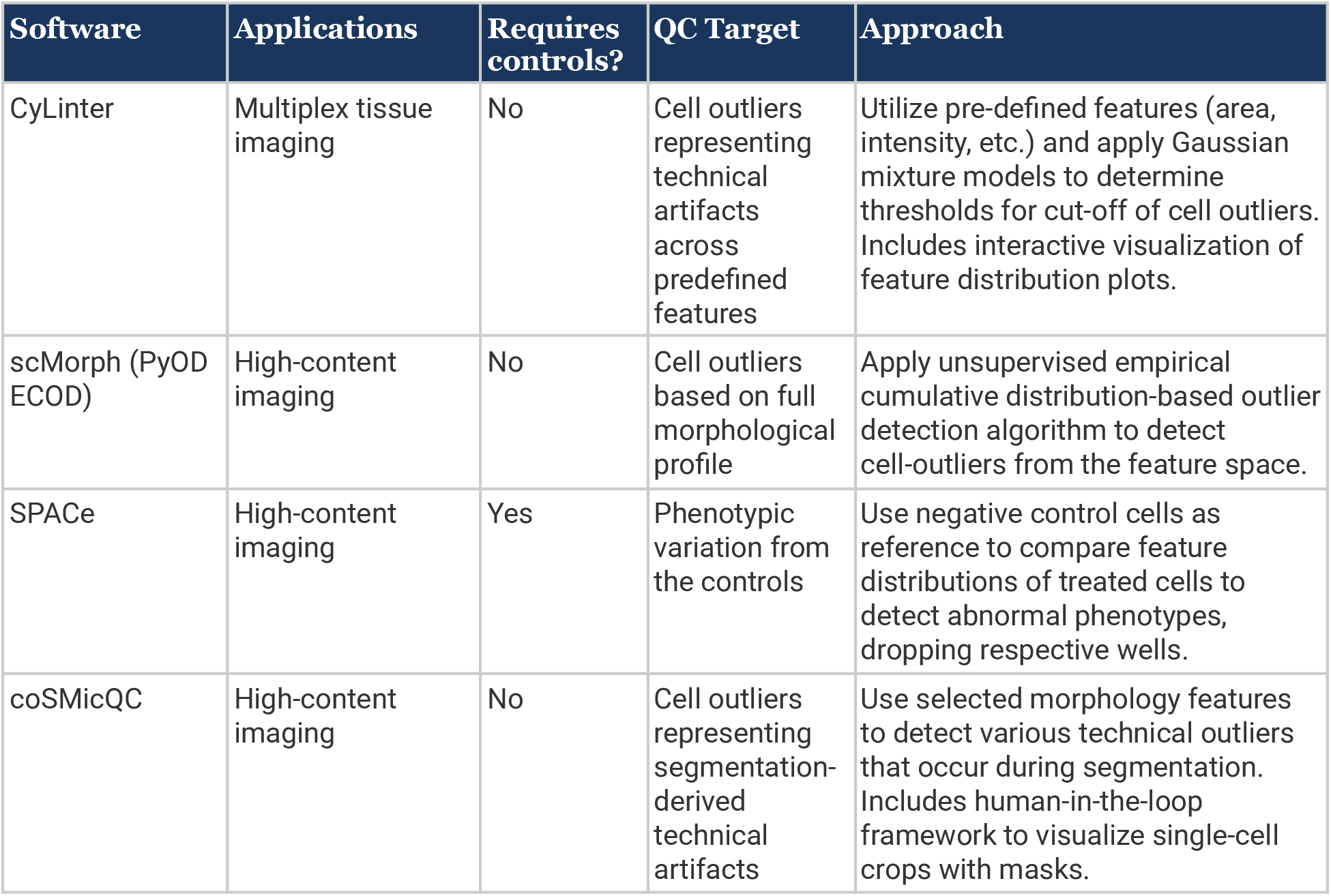
Overview of current open-source software with cell-level QC methods. This table compares four software tools that perform cell-level QC using various strategies. CoSMicQC focuses on detecting segmentation-derived outliers through the selection of a subset of features, distinguishing it from other software that broadly filters cells based on comparison to the controls or the entire feature space.

CoSMicQC works by distilling user-defined raw morphology features into what we call “conditions”. Conditions are descriptive of the specific technical issues that the underlying morphology features capture. For example, a condition may include detecting under-segmentation of nuclei. In the backend, coSMicQC z-score transforms raw feature values of the user-selected features, which converts the full feature vector into a mean of 0 and a standard deviation of 1. A coSMicQC user sets thresholds (in standard deviations) on these features to gate poor quality segmentations (**Supplemental Figure 1A**). Next, coSMicQC summarizes outlier cells, reporting the number of failed cells, proportion of failed cells per plate, and the feature range of the detected outliers for the condition features (**Supplemental Figure 1B**).

Another critical output of coSMicQC is a dataframe containing single-cell crops alongside feature values, which gives users the power of human-in-the-loop feedback to optimize condition thresholds. Behind the scenes, the dataframe is generated by additional software we developed, called CytoDataFrame.^39^ CytoDataFrame is an in-memory data model compatible with Jupyter notebooks (**Supplemental Figure 1C**). This rapid, human-in-the-loop interaction enhances user experience, reduces time managing disparate data sources, and we have found improves QC performance. Users can either filter or flag outlier cells before they continue with the image-based profiling pipeline. In practice, a user would filter low-quality single-cells to reduce noise in downstream analyses. A user can alternatively flag cells for downstream exploration (e.g., to assess the potential technical influence of outliers on downstream tasks).

### Optimizing assay conditions with coSMicQC

To illustrate coSMicQC in practice, we used it to optimize seeding density for cell lines that had never previously been Cell Painted. In theory, as cells become more crowded, segmentation becomes more challenging, so we expect more under-segmentation outliers, where multiple nuclei are merged into a single object. To test this, we analyzed one plate from a Cell Painting dataset spanning five cell lines, each measured at five different seeding densities at a 24-hour time point. We applied coSMicQC using two conditions based on nuclei features and one condition based on a cell feature to detect three kinds of segmentation artifacts (**Figure 2A**). To determine appropriate thresholds for each condition, we used an interactive Jupyter notebook environment that provides human-in-the-loop capabilities (via CytoDataFrame^39^). Thresholds represent the number of standard deviations (std) from the mean z-scored feature value and act as gates that distinguish poor-quality segmentations from real biology.

**Figure 2:**
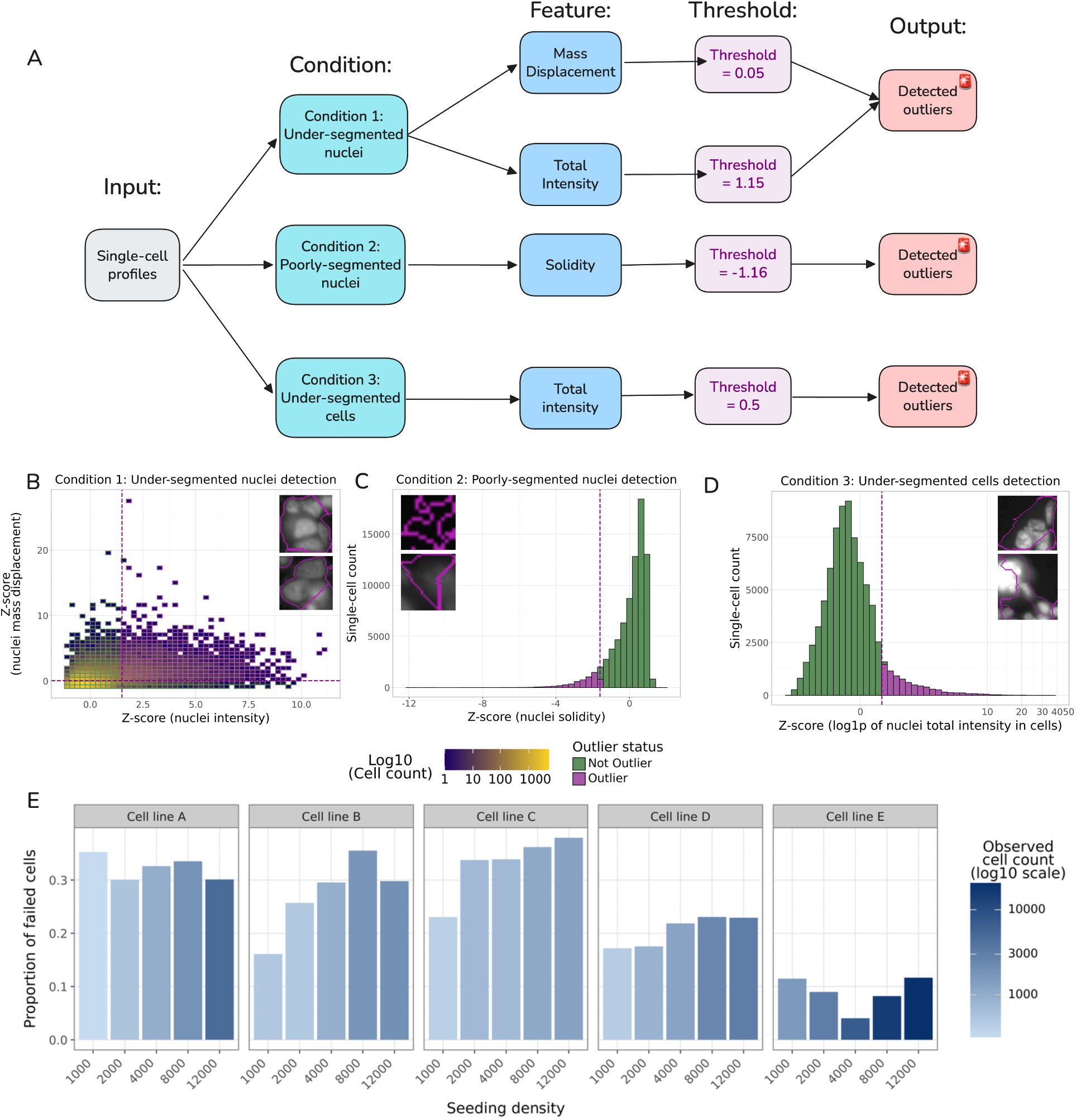
Example coSMicQC usage to identify poor quality segmentations. (**A**) A user interacts with coSMicQC to specify three “conditions” of different poor quality segmentations using four raw CellProfiler features. They then interactively set thresholds with real-time feedback of the single-cell images at the borderline regions of the threshold. In this example, a user identifies under-segmented nuclei, “poorly”-segmented nuclei (which can also include under-segmentations), and under-segmented cells. (**B**) Condition 1 thresholding to identify under-segmented nuclei caused by over-confluence of single-cells. (**C**) Condition 2 thresholding to identify “poorly” segmented nuclei, including the segmentation of non-nuclei background or over-segmentation. (**D**) Condition 3 thresholding to identify under-segmented cells that include nuclei of other cells. In both example conditions, we show randomly-selected example outlier single-cells with their overlaid segmentation outlines (magenta). (**E**) The proportion of cells failing QC across cell lines and seeding densities.The original cell lines’ names in order from A-E are as follows: CHLA-10, CHLA-113, CHLA-218, CHLA-25, and U2-OS.

#### (Condition 1) Detecting under-segmented nuclei (more than one nucleus within a single nuclei mask)

We used two CellProfiler features for this condition: mass displacement (the distance between the intensity-weighted centroid and the geometric centroid of the segmentation mask) and integrated intensity (the total Hoechst intensity within the nuclei). We set thresholds filtering segmented nuclei 0.05 std above the mean for mass displacement and 1.5 std above the mean for intensity (**Figure 2B**). When using two features within a single condition, we recommend setting separate thresholds for each feature.

#### (Condition 2) Detecting poorly segmented single nuclei (debris or over-segmentation; one nuclei split into multiple objects)

We used solidity, which quantifies indentations, or grooves, along the nuclei segmentation boundary. Low solidity indicates a highly non-circular object, and we expect nuclei, in most cases, to have a round, smooth shape. We set a threshold of 1.6 std below the mean for solidity to identify non-round and likely poorly segmented nuclei (**Figure 2C**).

#### (Condition 3) Detecting under-segmented cells

We used the integrated intensity of the Hoechst channel, which measures the total intensity of nuclear staining, within the cell compartment. We found that abnormally high values indicate cell segmentations that likely inappropriately contain multiple nuclei, which can occur when using CellProfiler (or other software that drops nuclei based on diameter size range, including QuPath^8^) for segmentation. Prior to cell segmentation, the segmentation module discards mis-segmented nuclei that are too large, based on user-specified cell diameter parameters. Removing these mis-segmented nuclei has a side effect that can cause under-segmentation of neighboring cells: the discarded regions, when they actually contain clumped nuclei, may still be incorporated into current cell segmentation masks, producing cell objects that do not accurately represent individual cells (see **Supplemental Figure 2**). We set a threshold of 0.5 std above the mean for nuclei-channel intensity within the cell compartment to identify under-segmentation of multiple cells into a single cell mask (**Figure 2D**).

We applied coSMicQC with these conditions and thresholds and observed an increase in the fraction of failing cells with increasing density in three of five cell lines (**Figure 2E**). At an FOV level, we observed a coSMicQC failure rate as high as ∼67%, driven by poor-quality conditions and clustering nuclei (**Supplemental Figure 3A**). Two cell lines (A and E) showed variable failure rates across seeding densities. For cell line A, this was consistent with our observation that lower densities caused clumping nuclei and mis-segmentations. Cell line E showed more instances of clustering or over-segmentation of nuclei at lower seeding densities, but proliferated rapidly at higher seeding densities (**Supplemental Figure 3B**). Taken together, these results show that coSMicQC conditions are robust to variability in cell morphology, density, and growth patterns arising from differences in cell behavior and culture conditions. This robustness allows us to use coSMicQC to select the seeding density that minimizes QC failures for each cell line.

We next compared coSMicQC to two manual annotators and observed that coSMicQC agreement was consistent with human-to-human agreement (**Supplemental Figure 4A**). The human annotators tended to classify more segmentations as poor quality, which resulted in coSMicQC showing lower and inconsistent sensitivity. We nevertheless observed high specificity and precision for coSMicQC, while the manual annotators had comparably low scores (**Supplemental Figure 4B**). Some cell lines were harder for coSMicQC to match with annotator labels (**Supplemental Figure 4C**), and harder for annotators to agree with each other (**Supplemental Figure 4D**), likely due to clumping cells (see **Supplemental Figure 3**). Overall, this analysis demonstrates that coSMicQC detects poor-quality segmentations with agreement comparable to manual annotators, while minimizing false positives, which would contaminate downstream analyses more than false negatives. In practice, coSMicQC users can toggle thresholds to balance false positives and false negatives.

### coSMicQC improves phenotype classification and outperforms PyOD

We applied coSMicQC to a Cell Painting dataset comprising two plates of cardiac fibroblasts. The first plate (“retransplantation plate”) contained cells from a healthy donor and from a patient with heart failure requiring retransplantation, treated with two different chemical perturbations plus a DMSO control. We used this plate to evaluate the impact of QC on clustering single-cell profiles. The second plate (“IDC plate”) contained cells from two healthy donors and four patients with idiopathic dilated cardiomyopathy (IDC).^28^ We used this plate to evaluate the impact of QC in predicting heart failure phenotypes. Because of the fibroblast growth patterns, we observed many instances of under-segmented nuclei and mis-segmented cells (**Supplemental Figure 5A**). We applied two coSMicQC “conditions” (different from the previous section) to both plates: one condition using nuclei area and nuclei total intensity to find mis-segmented nuclei, and the second condition using cell area to detect abnormally small, mis-segmented cells (**Supplemental Figure 5B**). CoSMicQC flags 19.3% and 19.2% of cells as poor-quality for the retransplantation and IDC plates respectively, with the majority detected as mis-segmented cells (**Supplemental Figure 5C**). We also compared coSMicQC to the Empirical Cumulative-distribution-based Outlier Detection (ECOD) algorithm from PyOD that the software scMorph^15^ uses for single-cell QC. PyOD contains a single parameter, the expected contamination fraction, which, to maximize comparison between methods, we set at 19.3% or 19.2% to match the proportion of cells flagged by coSMicQC.

We first analyzed the retransplantation plate. Both coSMicQC and PyOD ECOD removed 4,994 and 4,991 cells respectively out of the 25,859 observed cells, with only 47% of outliers shared. Transforming feature-selected single-cell morphology profiles with a Uniform Manifold Approximation and Projection (UMAP)^40^, we found that coSMicQC flagged cells that were largely isolated in one region of the UMAP space. In contrast, PyOD ECOD-detected outliers are more dispersed (**Figure 3A**). We visualized example single-cell segmentations and found that many PyOD ECOD outliers reflected genuine single-cell variation rather than segmentation artifacts compared to coSMicQC (**Supplemental Figure 6A**). After removing outliers, we re-fit UMAP and observed clusters corresponding to heart failure status and treatment combinations for both methods (**Figure 3B**). We sought to quantify these clustering improvements by applying Hierarchical Density-Based Spatial Clustering of Applications with Noise (HDBSCAN)^41^ to the single-cell profiles with or without QC from either method. We found that the feature space without QC applied (pre-QC) split into two clusters based on segmentation quality. After applying coSMicQC, the feature space split into three clusters that were separated based on biology (treatment and heart failure status) unlike PyOD ECOD (**Figure 3C**). To further compare QC methods, we calculated pairwise Pearson correlations between single cells in three groups: (1) cells that failed QC, (2) cells that passed QC, and (3) cells that failed QC compared to passing cells. Cells that fail coSMicQC have higher pairwise correlations compared to cells that fail PyOD ECOD, which form a bimodal distribution (Cohen’s d = 0.426, permuted p value = 0.0001). Furthermore, the pairwise correlation of passing and failing cells is higher for PyOD ECOD compared to coSMicQC (Cohen’s d = −0.208, permuted p value = 0.0001) (**Figure 3D**). Taken together, coSMicQC improved clustering of single-cell phenotypes and identified a more consistent population of poorly segmented cells than PyOD ECOD, which flagged more heterogeneous outliers, including a higher proportion of cells correlated with genuine biological variation.

**Figure 3:**
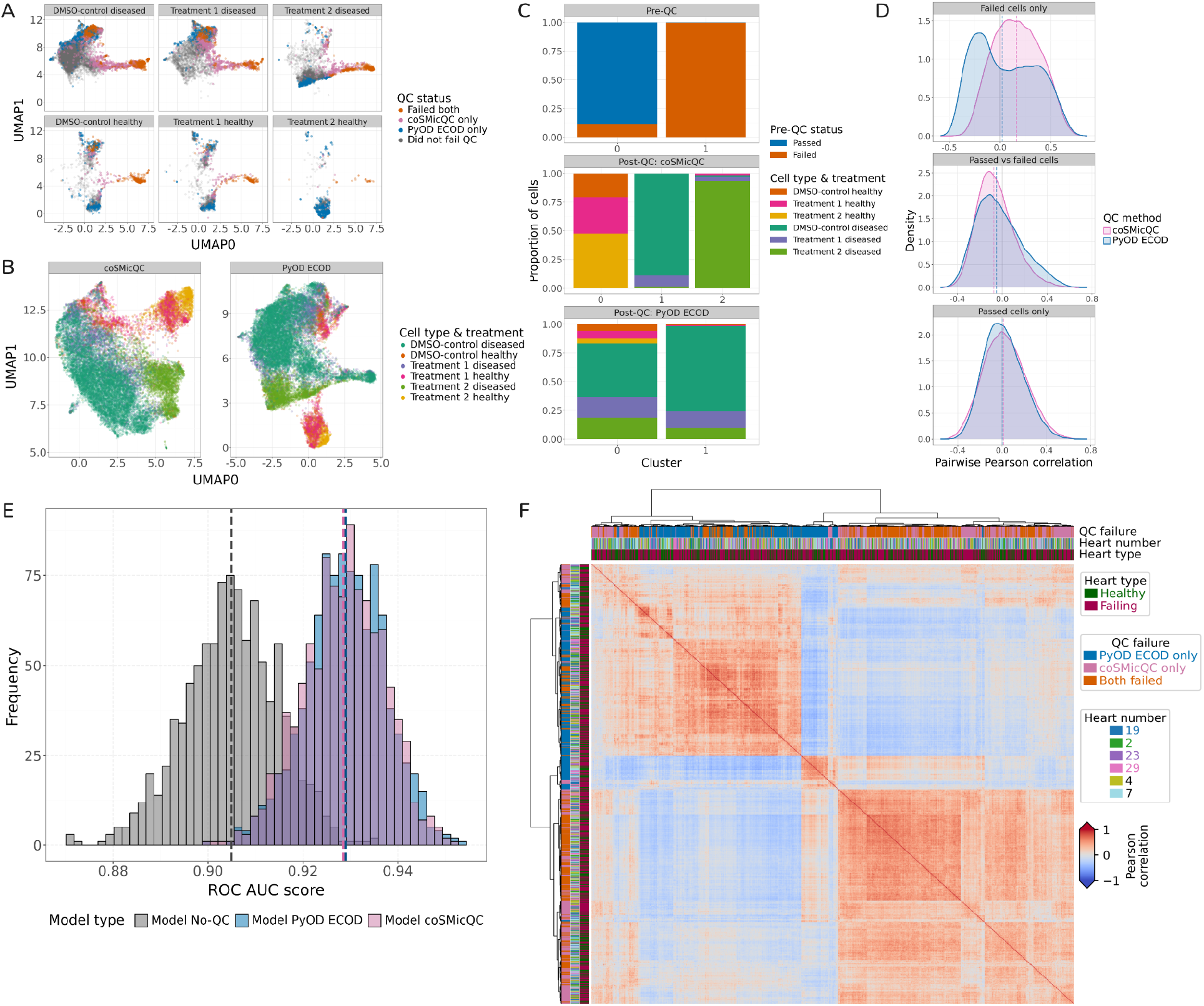
Single-cell QC with coSMicQC improves phenotype classification. (**A**) Transforming feature-selected, single-cell profiles with Uniform Manifold Approximation and Projection (UMAP) demonstrates that the large protrusion/cluster reflects poor quality segmentations. Outliers detected from coSMicQC overlap more with the protrusion than PyOD ECOD, which detects substantially more outliers in healthy phenotype. (**B**) After QC, the new UMAP transformation from coSMicQC suggests improved clustering while PyOD ECOD still shows an abnormal protrusion. (**C**) HDBSCAN confirms coSMicQC improves profile quality by identifying three clusters based on biological phenotypes (cell type and treatment). (**D**) Pairwise Pearson correlations between cells that failed coSMicQC are substantially higher than PyOD ECOD, indicating that coSMicQC consistently detects segmentation artifacts that will look morphology similar. PyOD ECOD demonstrates a bimodal distribution of correlations between failed cells, indicating more variable outliers that could include biological phenotypes. PyOD ECOD also has higher correlations between failing and passing cells, indicating that there is a similar biological phenotype within the detected outliers. The dotted lines reflect the median for the distributions. (**E**) Machine learning phenotype classification performance significantly improves after applying coSMicQC, with very similar performance between PyOD ECOD and coSMicQC. (**F**) Cells that failed coSMicQC only are more similar to cells that failed both QC methods than PyOD ECOD, indicating consistency in detected outliers.

We next analyzed the IDC plate to measure the impact of QC on phenotype signature discovery. Both coSMicQC and PyOD removed 3,996 and 4,005 cells respectively out of 20,856 observed cells, with only a 54% overlap. We trained three logistic regression models (one on pre-QC profiles and one on post-QC profiles, per QC method), to predict whether a cardiac fibroblast came from a healthy donor heart or a heart from a patient with IDC. Similar to Travers et al, we held out the DMSO-treated wells from one healthy heart and one failing heart.^28^ We then applied each trained model to its corresponding holdout wells, generating predicted classes and probabilities for each holdout cell. To estimate variability in model performance, we bootstrapped single cells in the holdout wells 1,000 times per QC method. For each iteration, we resampled about 20% of cells without replacement and calculated the Receiver Operating Characteristic Area Under the Curve (ROC AUC) (see **Methods** for details). We observed a mean ROC AUC increase of 0.024 for coSMicQC (95% CI: 0.000 to 0.049), outperforming pre-QC in 96.6% of bootstrap comparisons (**Figure 3E**). We observed slightly higher performance for PyOD ECOD (**Supplemental Figure 6B**). We next calculated pairwise Pearson correlations between all cells that failed QC by either method. PyOD ECOD only failures (blue) formed a distinct cluster separate from coSMicQC only failures (pink), and cells that failed both methods (orange) correlated more strongly with cells that failed only coSMicQC (**Figure 3F**). We next sought to compare the impact of QC on machine learning model performance, by applying the same three models to cells that failed QC by either method. The pre-QC model showed high confidence for some failed-QC cells in both classes (healthy and diseased), suggesting either that many cells were inappropriately filtered by QC, or that models trained on technical outliers are biased by those outliers. The post-QC models, when applied to these same failed-QC cells, also showed high, though somewhat dampened, confidence scores, indicating incomplete and uneven QC filtering across phenotypes (**Supplemental Figure 6C**). Taken together, analyses from both plates show that QC of any kind improves single-cell phenotype discovery, but coSMicQC filters a higher proportion of technical outliers relative to biological signal than PyOD ECOD. Our data also suggest that technical bias can creep into a dataset in ways QC minimizes but cannot always fully remove. As with any experiment, we recommend careful experimental design, such as optimizing plating conditions and segmentation parameters, as the first line of defense for reliable biological signal detection.

### coSMicQC identifies cell culture contamination

We developed a coSMicQC function to detect abnormal perinuclear signals, which can represent cell contamination (e.g., mycoplasma) and high channel overlap or bleedthrough. Because nuclear stains bind DNA confined in the nuclear region, we do not expect signals outside the nucleus. We used CellProfiler measurements of the nuclear channel for detecting abnormal perinuclear signal.^42^ The coSMicQC perinuclear signal detector works in a three-step procedure (**Supplemental Figure 7A**).

#### Step 1: Determine if there is an abnormal signal present

The coSMicQC perinuclear signal detector calculates two measurements: (Measurement 1) *Skewness of cytoplasm texture around the nuclei* and (Measurement 2) *Nuclear form factor variability*. We derive measurement 1 by calculating Bowley’s skewness score^43^ of the CellProfiler feature “Information Measure of Correlation 1” in the nuclear channel measured in the cytoplasm compartment. Bowley’s skewness flags an unexpected, non-Gaussian distribution for this CellProfiler feature, which we observed occurs when cells have abnormal perinuclear signals (**Supplemental Figure 10**).

Based on empirical observations, we set skewness thresholds at −0.15 and 0.09, capturing both left- and right-skewed distributions. We derive measurement 2 by calculating the interquartile range (IQR) of the CellProfiler feature “Form Factor” in the nuclei compartment. We expect the vast majority of nuclei to be round in shape, and this measurement flags an abnormally non-circular population of nuclei. Based on empirical observations, we set the IQR threshold at an upper bound of 0.15. If both measurements pass the thresholds, then we consider the plate clean of abnormal perinuclear signals. If either of the measurements fails, then we move on to step 2. Our approach was to set reasonable default thresholds based on empirical evidence, which we carry over to future projects. A user can update these thresholds if necessary, but we do not recommend this advanced usage.

#### Step 2: Determine if an abnormal signal occurs in the whole plate

The conventional “condition” system in coSMicQC relies on standard deviations calculated from distributions, which will fail if an entire plate is compromised. The condition system will fail because coSMicQC runs per plate, and if the entire plate suffers from abnormal perinuclear signal, all the cells will be poorly-segmented and thus shift entire feature distributions. To account for this, we calculate the mean of the two raw CellProfiler nuclei features used in the previous step. We consider the full plate compromised if coSMicQC measures a high mean texture value greater than −0.25 (corresponding to abnormally heterogeneous pixel intensity) or a low mean nuclear form factor value lower than 0.78 (indicating a high proportion of non-circular nuclei). We display the outcomes of these calculations in a 2×2 decision table (see **Supplemental Figure 8A**).

Specifically, if both raw feature means exceed thresholds, then coSMicQC detects abnormal signal across the whole plate and the procedure terminates. If both raw feature means do not exceed thresholds, then coSMicQC detects partial abnormal signals on the plate, and we move on to step 3. If only the nucleus Form Factor measurement exceeds thresholds, then coSMicQC will recommend verifying appropriate segmentation parameters.

#### Step 3: Detect which wells are compromised

If the coSMicQC perinuclear signal detector arrives at step 3, then we use the standard coSMicQC outlier detection approach, setting two conditions. The first condition uses cytoplasm texture around the nuclei (the same raw measurement in Step 2), and the second condition uses cytoplasm granularity in the nuclei channel (**Supplemental Figure 8B**). For the second condition, we specifically use the CellProfiler feature cytoplasm granularity at the second structuring element size in the nucleus channel, which describes fine speckles or small structures around the nucleus that are common in cells that have abnormal perinuclear signals. At step 3, a user will interact with CytoDataFrame to set dataset-specific thresholds for compromised wells, and it will report which wells have the highest proportion of compromised cells.

We applied coSMicQC’s perinuclear signal detector to a dataset containing three different Schwann cell lines.^44^ We previously observed mycoplasma contamination in one of these cell lines (**Figure 4A**). The coSMicQC perinuclear signal detector arrived at step 3 and clearly distinguished cells with contamination (**Supplemental Figure 8B**), flagging a substantially higher proportion of contaminated cells across wells of the contaminated cell line (**Figure 4B**). Wells in other cell lines show minimal flagged cells, which we concluded, after visual inspection, were due to minor channel overlap (channel bleedthrough) in the nuclear channel, as expected to be potentially flagged as abnormal perinuclear signal. Using the embedded CytoDataFrame functionality, we observed that the flagged cells show heavy mycoplasma contamination in the nuclei channel (**Figure 4C**).

**Figure 4:**
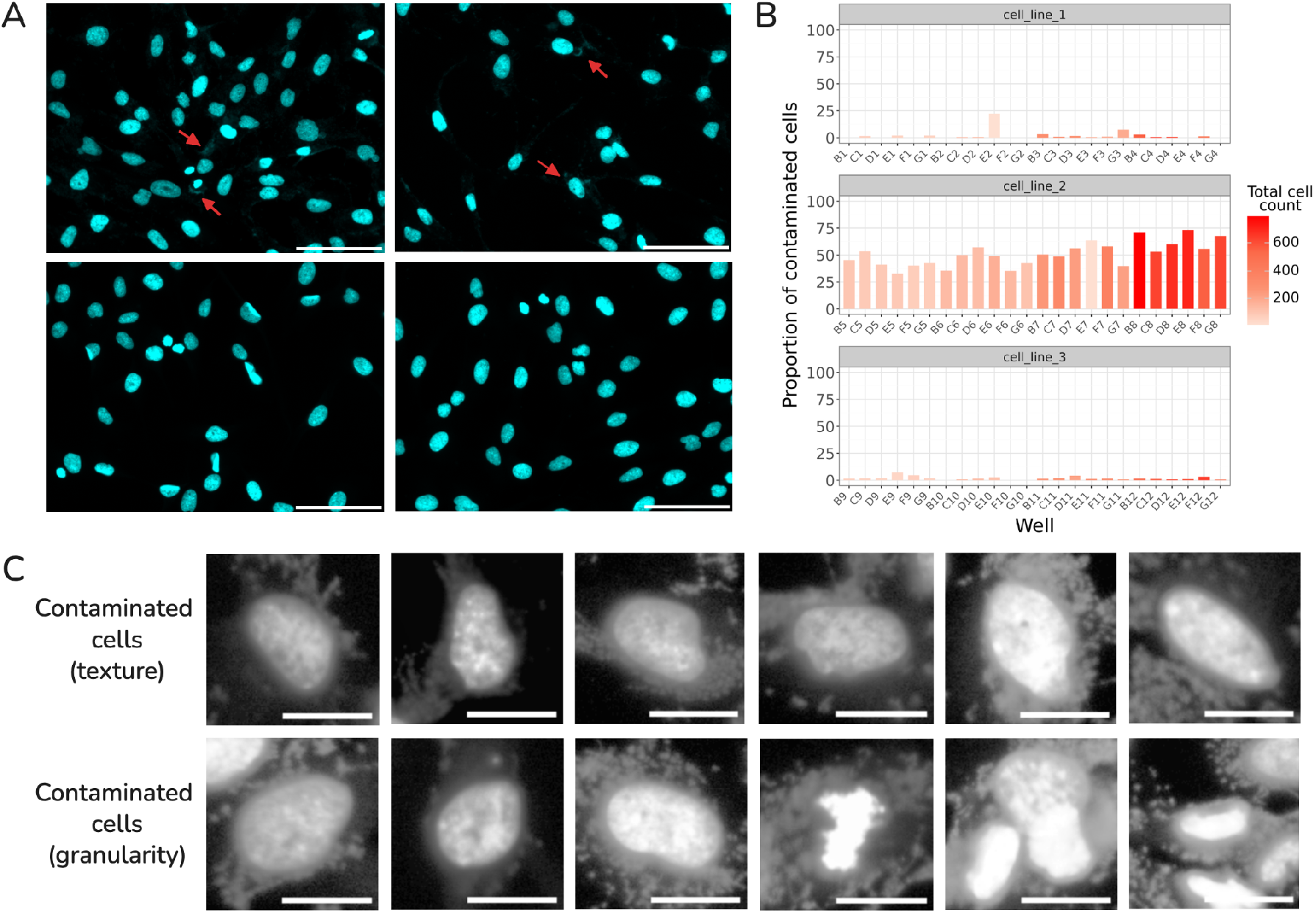
The coSMicQC perinuclear signal detector flags mycoplasma contamination. (**A**) Field of view (FOV) examples show Schwann cells either contaminated with mycoplasma (top) or uncontaminated (bottom). Red arrows highlight visible mycoplasma for these cells. The scale bar is 100 μM. (**B**) The proportion of single-cell QC failures per well shows a single cell line with a high proportion of flagged contaminated cells compared to other cell lines. (**C**) Example single-cell crops from the CytoDataFrame Jupyter notebook viewer within coSMicQC show single cells with mycoplasma contamination. The coSMicQC detector uses two conditions to detect abnormal perinuclear signals (see **Supplemental Figure 7**). The top row shows detected compromised cells using cytoplasm texture around the nuclei, and the bottom row shows compromised cells detected from cytoplasm granularity around the nuclei. The scale bar is 20 μM. The Schwann cells from each genotype were derived from the same ipn02.3 2λ cell line, and the original names from 1-3 are as follows: NF1 wild-type genotype (A3), NF1 heterozygous genotype, and NF1 null genotype (C04).

### coSMicQC has little impact on MOA prediction but does alter lead compound ranks in a high-throughput drug screen

We reprocessed the LINCS Cell Painting dataset^17^ using CytoTable for single-cell profile harmonization, coSMicQC for single-cell QC, and Pycytominer for aggregation, normalization, and feature selection. We applied the same quality control conditions and thresholds as in Figure 2. We first reproduced the UMAP figure from the original paper, which did not apply single-cell QC (**Figure 5A**). We then applied QC, and observed a concentrated region in UMAP space that contained a high proportion of cells per well that failed QC (**Figure 5B**). We removed the failed single cells from these profiles, reprocessed data through the image-based profiling pipeline, and recalculated UMAP coordinates. We observed a restructuring of the UMAP space, with more condensed clusters and an exclusion of many of the small, isolated clusters that were in the original UMAP (**Figure 5C**). Next, following an approach similar to the original LINCS paper (see **Methods** for details), we trained a multi-class neural network to predict compound MOA and compared performance with and without QC. We see substantial performance increases for the three MOAs that improved most after QC, and similarly substantial decreases for the three MOAs that declined most after QC (**Supplemental Figure 9A-B**). However, we observed only a small effect of QC overall, with 25/222 MOAs (11%) showing a change in performance from the test set, suggesting that the aggregation procedure largely dampens the impact of QC on MOA prediction (**Supplemental Figure 9C**). CoSMicQC flags cells at a similar rate regardless of whether an MOA’s prediction performance improved or declined after QC, indicating no overall meaningful change to MOA prediction performance (**Supplemental Figure 9D**). For traditional, aggregation-based applications of image-based profiling, coSMicQC has a comparatively modest impact.

**Figure 5:**
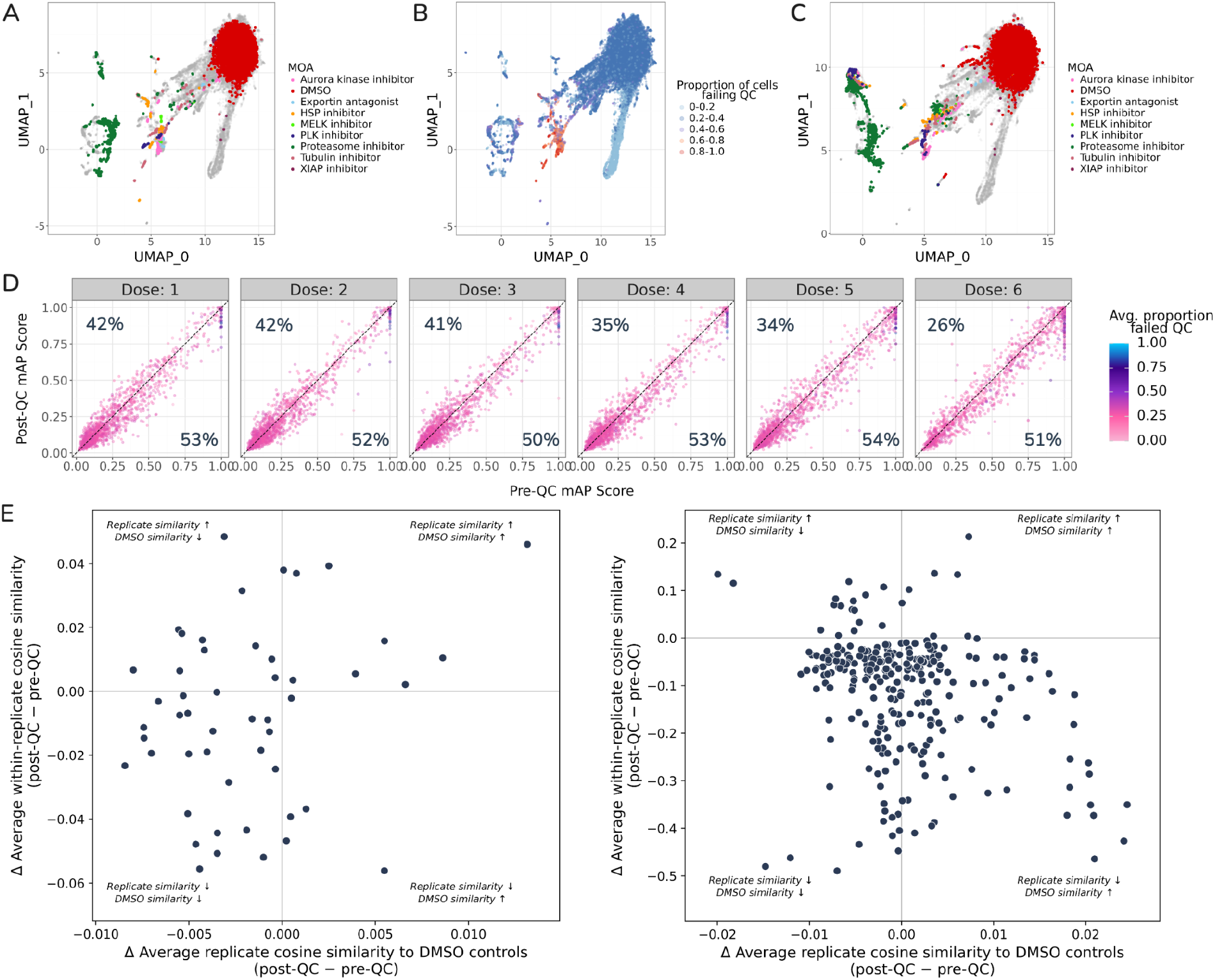
coSMicQC impacts UMAP clustering and lead compound nomination. (**A**) Reproduced UMAP from the original LINCS Cell Painting paper (Way et al. 2022), where no QC was applied, showing many isolated and loosely coupled clusters. (**B**) UMAP of morphology features without QC, colored by the fraction of segmented cells classified as QC failures within each well. (**C**) UMAP of the LINCS dataset after applying single-cell QC with coSMicQC reduces isolated clusters. Across doses, coSMicQC **(D)** decreases mAP for some compounds and increases it for others. **(E)** CoSMicQC impacts lead compound nomination, comparing (left) compounds that were rescued and (right) compounds that were demoted in rank after quality control. Most rescued compound-dose pairs show decreased similarity to DMSO controls, with replicate similarity declining at a lower rate than for compounds whose mAP decreased post-QC.

We next calculated mean average precision (mAP)^45^ scores for compounds without QC (pre-QC) and with QC (post-QC) across all doses separately. mAP is a metric that captures replicate reproducibility and phenotypic difference from controls, where a high value indicates a treatment that is both reproducible and phenotypically distinct from controls. We observed, on average across doses, a reduction in mAP in the post-QC dataset for 52% of compounds and an increase for 37% of compounds indicating a variable effect of coSMicQC on compound reproducibility (**Figure 5D**). 11% of compounds across doses showed no change. This was an unexpected finding, as we had anticipated that coSMicQC would improve mAP for the majority of compounds by revealing more underlying biology. We investigated further, examining the relationship between the fraction and nature of cells failing QC and the resulting change in mAP post-QC. We observed a negative relationship between the fraction of cells removed and post-QC mAP: compounds with a high fraction of cells removed showed the largest drops in mAP (**Supplemental Figure 10A**). Calculating cosine similarity for replicates post-QC, we found that compounds with decreased mAP had significantly lower cosine similarities than compounds with increased mAP (**Supplemental Figure 10B**). Finally, we calculated mAP using only cells that failed QC, and we observed significantly higher scores than when using only cells that passed QC suggesting that compounds with decreased mAP post-QC had been artificially inflated pre-QC by poor-quality cells, which resemble each other more closely than the true heterogeneity of genuine cells (**Supplemental Figure 10C**). We next ranked compounds by mAP score both pre- and post-QC, with a score of 1 corresponding to rank 1 and tied scores sharing the same rank (rank method=’min’). QC affected many compounds in the top-20 rankings across all doses (**Supplemental Figure 11A**). In total, coSMicQC nominated 50 treatments into a new top-20 ranking per dose: 43 unique compounds rescued at one dose, two unique compounds rescued at two doses, and three unique compounds reduced at three doses (**Supplemental Figure 11B**; across five doses, there are 100 total top-20 spots). Surprisingly, QC did not significantly increase pairwise cosine similarity among replicates for the rescued compounds (**Supplemental Figure 12)**. Instead, their migration into the top 20 reflected a combination of two factors: these compounds became less similar to DMSO controls, while their replicate similarity declined at a lower rate than it did for compounds whose mAP decreased post-QC (**Figure 5E**). Taken together, these findings tell a nuanced story. Compounds with high QC failure rates appear to have inflated pre-QC scores, because their segmentation errors tend to resemble one another and inflate apparent replicate reproducibility. At the same time, removing technical outliers from the DMSO controls allowed genuine biological differences to emerge. Removing these low-quality cells and artifacts therefore yields a more biologically grounded evaluation of these compounds, even in a single-cell aggregation regime.

## Discussion

CoSMicQC is a reproducible Python package that users can install via traditional channels (e.g., PyPI). We document coSMicQC functionality through multiple example notebooks, blog posts, and documentation. The software fits seamlessly with the traditional image-based profiling pipeline. CoSMicQC leverages a simple Pythonic approach and only requires minor configuration tweaks. It filters or optionally flags single-cell technical outliers based on user-defined conditions representing different categories of poor segmentations (e.g., over-segmentation or incorrectly segmenting debris). For example, if nuclei were under-segmented, we tend to see more non-smooth segmentations (high quantity of grooves) in masks where the algorithm did not outline objects correctly. A measurement known as “nuclear solidity”, which ranges from 0 to 1, where 1 is a perfect circle, helps objectively describe this detail. Multiple features can be used together to identify technical outliers, such as nuclear area and mass displacement for overlapping nuclei segmentations. We engage human-in-the-loop workflows in coSMicQC through CytoDataFrame, which dynamically visualizes single-cell crops with optionally overlaying segmentation outlines to verify threshold configuration and single-cell technical outliers.

We demonstrated the ability of coSMicQC to detect technical outliers and showed how single-cell QC improves image-based profiling. When applied to single-cell applications, coSMicQC optimized assay conditions, improved clustering after removing poor-quality single cells, improved performance in single-cell phenotype prediction, and outperformed modern outlier algorithms in maximizing detection of segmentation artifacts and minimizing filtering rare biological phenotypes. When applied as a preprocessing step before aggregation in a high-throughput drug screen, coSMicQC altered lead compound nomination but did not impact supervised MOA prediction. We demonstrate that. Adding coSMicQC to either single-cell or aggregated image-based profiling pipelines is likely to improve data quality and thus the insights gained from these experiments. Furthermore, we demonstrated how coSMicQC can accurately detect abnormal perinuclear signals using nuclear morphology features. Mycoplasma contamination in cell cultures can happen across laboratories^46^ and is not always detectable by eye.^47,48^ Mycoplasma alters cell morphology^49,50^, which makes detection important prior to downstream analysis and machine learning. A user should use the perinuclear signal detector as a backup plan. Other forms of technical issues can also occur as abnormal perinuclear signals, such as channel overlap or spectral bleed through where other cell compartments can be seen in the nuclear channel. If this flag is triggered, we recommend reviewing flagged images or the entire plates carefully. A limitation of using the nuclear channel and features to detect abnormal perinuclear signals is that other forms of contamination may not be flagged.

Another limitation is that the detector by default expects CellProfiler features, and a user may need to reconfigure thresholds per dataset. We focused on nuclear features because existing mycoplasma detection methods use DAPI (or Hoechst) staining.^42^ The standard Cell Painting protocol also uses DAPI staining, which simplifies methodology and expands use cases to detect both potential contamination and significant channel overlap. We evaluated coSMicQC using Cell Painting datasets, but it can be applied to any high-content microscopy assay that applies single-cell segmentation and extracts interpretable, single-cell morphology features (e.g., from CellProfiler).

Another limitation of coSMicQC is that its z-score threshold approach requires a sufficient baseline of good-quality cells to accurately flag segmentation errors. CytoDataFrame addresses this by letting users visualize cells in real time to detect plate-wide quality issues. More broadly, coSMicQC risks removing false positives, cells flagged as artifacts that may in fact be biologically relevant. Overzealous thresholds therefore risk removing genuine biology along with artifacts, reducing discovery potential. coSMicQC mitigates this risk through the human-in-the-loop feedback enabled by CytoDataFrame, allowing users to dynamically set thresholds based on their specific hypotheses or dataset. PyOD ECOD shares this limitation, since it likewise requires a contamination fraction parameter set by the user. The goal of coSMicQC and single-cell QC in general is to minimize the removal of interesting phenotypes while maximizing the removal of technical outliers. Currently, coSMicQC identifies outliers using z-score-based thresholds defined in terms of standard deviations from the mean for a subset of features. As a result, the underlying feature distributions may vary across experiments, requiring users to evaluate and, if necessary, adjust thresholds (or generate new conditions) for each plate. We recognize that this trade-off is never perfect in practice, but coSMicQC gives users control over how to balance it. As the community gains experience with the tool, we expect more robust default thresholds and more comprehensive detection conditions to emerge, reducing the need for manual fine-tuning.

We evaluated coSMicQC using traditional segmentation methods from CellProfiler, which includes three modules with manual parameters to optimize segmentation of nuclei and cells. State-of-the-art (SOTA) segmentation methods, such as those based on the Segment Anything Model (SAM) architecture, improves segmentation accuracy when applied to fluorescence microscopy datasets.^51–53^ For example, a recent study comparing segmentation algorithms showed SAM outperforms Cellpose v3 (cyto model).^54^ However, this study only evaluated performance using one dataset and the well-characterized and easy-to-segment A549 cell line (in optimal conditions). In a more comprehensive benchmark, Achit and Page 2026 evaluated eight SAM-based segmentation methods across 36 image sets from various imaging modalities, including fluorescence microscopy.^55^ Across datasets, performance varied substantially, with high performance on another easy-to-segment U2OS cell line dataset, while demonstrating low performance on other more challenging cells. Overall, these results show that poor quality segmentations arise even with SOTA models when applied to challenging or heterogeneous high-content screens, leading to a need for QC methodologies to reduce the burden of poor quality segmentations on downstream analyses. Furthermore, coSMicQC requires computer vision features, such as those extracted by CellProfiler, and CytoTable harmonization. It is not restricted to CellProfiler outputs; any software or pipeline that exports profiles with metadata and computer vision features can be used, including scikit-image^56^, Squidpy^57^, and cpmeasure^58^. In other words, coSMicQC requires an interpretable feature space, which excludes deep learning methods such as DeepProfiler^37^). It also benefits from baseline knowledge of what each extracted morphology feature represents, which prepares users to detect segmentation outliers and interpret results downstream, but requires additional time and effort to learn and optimize thresholds for each dataset.

CoSMicQC is a reproducible home for single-cell QC in image-based profiling. By providing a reliable tool, we hope to persuade the field toward greater caution in data processing as it evolves toward single-cell analysis. Manually reviewing all FOVs to ensure accurate single-cell segmentation is nearly impossible, especially in high-throughput screens. Scientists instead rely on time-intensive manual inspection of a few representative FOVs per well to detect segmentation issues, an important process that is nevertheless subjective, narrow, and irreproducible. coSMicQC reduces this burden by programmatically detecting technical outliers without sacrificing human-in-the-loop decision making. Through this work, we present a single-cell QC standard that can be applied across pipelines to move the field toward the highest-quality image-based profiling.

## Methods

### Developing coSMicQC software

We developed coSMiCQC using Python, and we maintain compatibility with major supported versions. CoSMiCQC software is open-source and hosted on GitHub^36^. We manage version control with Git, and we distribute coSMicQC via PyPI for easy installation. We created the documentation with Sphinx, which includes automated builds, tutorials, and example pipelines.

CoSMicQC depends on standard scientific Python libraries, including NumPy^59^, pandas^60,61^, scikit-image^56^, and scikit-learn^62^ for the detection of outliers and contamination. The software also uses matplotlib^63^ and seaborn^64^ for visualization, specifically for the perinuclear signal detector when plotting the proportion of failed cells across wells. All dependencies, including development dependencies, are specified in the ‘pyproject.toml’ available in the repository.

Our sustainability practices include using pre-commit to ensure consistent code formatting and style. We utilize the Software Gardening Almanack^65^ as a pre-commit hook to perform linting checks that promote sustainable practices within the repository. We perform automated tests through GitHub Actions to verify functionality and cross-platform compatibility before updates are pushed. At the time of publishing coSMicQC has 95.1% code coverage.

### Using coSMicQC to optimize assays

For showcasing coSMicQC functionality in Figure 2, we used an unpublished Cell Painting dataset^66^ to analyze one plate of five pediatric cancer cell lines profiled at five different seeding densities and a 24-hour incubation time. We derived single-cell image-based profiles from CellProfiler using the standard segmentation modules (e.g., IdentifyPrimaryObjects, IdentifySecondaryObjects, IdentifyTeritaryObjects with manually optimized parameters) deriving nuclei, cell, and cytoplasm segmentations. CellProfiler segmentation and feature extraction pipeline can be found on Zenodo.^66^ The original cell lines’ names in order from A-E are as follows: CHLA-10, CHLA-113, CHLA-218, CHLA-25, and U2-OS. We applied coSMicQC as a scoring metric to identify most optimal conditions for each cell line independently. We used three conditions to flag under-segmented nuclei, poorly or mis-segmented nuclei, and under-segmented cells, and applied them to the entire plate.

#### Comparing coSMicQC to manual annotation

To build a dataset comparing coSMicQC labels to manual annotators, we first categorized cells into ten groups, either failing or passing QC based on coSMicQC labels and cell lines (2 QC statuses X 5 cell lines = 10 groups). We randomly selected 40 cells from each group (400 cells total) and provided the same randomly sampled cells to two human annotators. We asked them to manually annotate each cell as either a “good” segmentation, a “bad” segmentation, or “not sure” in case the single-cell crop is not clear enough to make a decision. We dropped cells with “not sure” annotations for either human annotator. We used CytoDataFrame and the computed CellProfiler bounding boxes to visualize the single-cell for the manual annotators (link to notebook: https://github.com/cytomining/coSMicQC/blob/main/docs/manuscripts/cosmicqc_paper_v1/supp_figure_x_alsf_manual_annotations/0.perform_manual_annotations.ipynb. We computed Cohen’s kappa agreement score between three groups (1) between human annotators, (2) between coSMicQC labels and annotator 1, and (3) between coSMicQC labels and annotator 2. We calculated precision, accuracy, sensitivity, specificity, and confusion matrices for each group considering the manual annotator to be ground truth and failing cells to be positive predictions. For comparing human annotators, we used annotator 1 as ground truth.

### Improving phenotype prediction with cosMicQC

We used the publicly-available dataset from Travers et al.^28^ The dataset contains two separate plates. The first plate, known as the “retransplantation” plate, contains two patient hearts (one failing after a retransplantation and non-failing) with two compound treatments and DMSO control. Treatment 1 refers to a transforming growth factor beta receptor inhibitor and treatment 2 is blinded with the alias of “drug_x”. The second plate, known as the “idiopathic dilated cardiomyopathy” (IDC) plate, contains cardiac fibroblasts from six patient hearts (two non-failing and four failing with IDC) with no treatments (media-only). We use the retransplantation plate exclusively to evaluate how QC improves recovery of biologically relevant signals in the presence of perturbations and controls, while we use the IDC plate to train models with and without QC to predict heart failure status under baseline conditions and evaluate changes in performance after QC. We acquired the morphology profiles using the default CellProfiler segmentation modules (e.g., IdentifyPrimaryObjects, IdentifySecondaryObjects, IdentifyTeritaryObjects with manually optimized parameters) deriving nuclei, cell, and cytoplasm segmentations. We previously normalized extracted CellProfiler features using Pycytominer z-score normalization (See Travers et al. for more details). CellProfiler segmentation and feature extraction pipeline can be found on Zenodo.^28^

#### Evaluating PyOD and coSMicQC in single-cell phenotype clustering

For the retransplantation plate, we generated three UMAP embedding spaces (pre-QC, coSMicQC applied, and PyOD ECOD applied) computed from respective feature-selected profiles. To identify groups of cells within the retransplantation plate feature spaces (pre-QC, coSMicQC, and PyOD ECOD), we performed principal component analysis (PCA) and selected the top five principal components, representing 36.99% variance for the pre-QC profile, 32.96% variance for the profile with coSMicQC, and 33.91% for the profile with PyOD ECOD, for downstream analysis. We applied HDBSCAN^41^ to the PCA-transformed feature spaces using a minimum cluster size of 50.

We calculated pairwise Pearson correlations using feature-selected profiles in three ways: (1) cells that failed QC, (2) cells that passed QC, and (3) cells that failed QC compared to passing cells. To compare failing vs. passing cells, we randomly sampled the same number of cells to create equally sized groups, which resulted in 50,000 randomly sampled pairwise comparisons per comparison group. For each QC method, we computed Cohen’s d comparing the resulting within-method correlation distributions between cells that failed QC versus cells that passed QC. To directly compare QC methods, we quantified the difference between the coSMicQC and PyOD ECOD cross-group correlation (passing versus failing cells) distributions using Cohen’s d. We assessed statistical significance using a permutation test with 10,000 permutations, in which we independently randomized QC failure labels per QC method while preserving the observed number of failed cells. We used the resulting null distributions to calculate a two-sided permutation p-value.

#### Evaluating PyOD and coSMicQC in machine learning predictions

For the IDC plate, we trained two binary logistic regression models using either coSMicQC or the PyOD ECOD QC method. We applied the same model hyperparameters, training/testing split, and holdout-heart evaluation strategy in both cases. We then performed bootstrapping by resampling 20% of cells from the pre-QC held out dataset (799 cells) and matching that same sample size for both QC held out sets, without replacement for 1,000 iterations, selecting both their true labels and predicted probabilities for each resample. We computed the ROC AUC for each resampled dataset, producing a distribution of scores pre- and post-QC. We computed the mean difference of the ROC AUC scores across bootstraps pre-QC and post-QC and reported the 95% confidence interval. In addition, we calculated the empirical probability of the post-QC model outperforming the pre-QC model by computing the proportion of bootstrap comparisons in which the post-QC model had a higher score than the pre-QC model across resamples. We split the cells that failed QC into three groups 1) failed coSMicQC only, 2) failed PyOD ECOD only, and 3) failed both methods. We randomly sampled 20% of cells from each group and computed pairwise Pearson correlations.

### Perinuclear signal detection with coSMicQC

We used the publicly-available dataset from Tomkinson et al.^40^, which includes CytoTable harmonized single-cell profiles derived from CellProfiler SQLite outputs. We acquired the morphology profiles using default CellProfiler segmentation modules (e.g., IdentifyPrimaryObjects, IdentifySecondaryObjects, IdentifyTeritaryObjects with manually optimized parameters) deriving nuclei, cell, and cytoplasm segmentations. CellProfiler segmentation and feature extraction pipeline can be found on Zenodo.^67^ We applied the coSMicQC perinuclear signal detector to one plate from the dataset. We confirmed by visual inspection that the cell line that was programmatically detected as contaminated contained a Mycoplasma infection. The Schwann cells from each genotype were derived from the same ipn02.3 2λ cell line, and the original names from 1-3 are as follows: *NF1* wild-type genotype, *NF1* heterozygous genotype, and *NF1* null genotype.

### Applying coSMicQC in an aggregation pipeline impacts lead compound rankings

We used the publicly-available dataset used Way et al.^17^ from the Library of Integrated Network-Based Cellular Signatures (LINCS) Cell Painting dataset, which is accessible via the Cell Painting Gallery^60^ under the dataset ID “cpg0004-lincs”. The project contains 136 plates, with 1,571 compounds measured across 6 doses in the A549 cell line. The original authors acquired the morphology profiles using CellProfiler default segmentation modules (e.g., IdentifyPrimaryObjects, IdentifySecondaryObjects, IdentifyTeritaryObjects with manually optimized parameters) to segment nuclei, cells, and cytoplasm. CellProfiler segmentation and feature extraction pipeline can be found on GitHub at this link: https://github.com/broadinstitute/imaging-platform-pipelines/tree/3eb4ff5676aa7889666f09b606cd915c8b9ea839/cellpainting_a549_20x_phenix_bin1

#### Reproducing LINCS image-based profiling

We applied CytoTable to the CellProfiler SQLite outputs to harmonize features for single-cell profiling. We then applied coSMicQC to detect under-segmented nuclei, over or mis-segmented nuclei, and under-segmented cells (containing multiple nuclei) using the same conditions and thresholds as applied in Figure 2. Cells failing these conditions were flagged in the profiles. We applied the same Pycytominer pipeline for aggregate, annotate, and normalize to DMSO controls as performed in the original paper. We also applied the same spherize and feature selection pipeline. The original paper recoded treatment doses as integer labels, which we also use here (“1” = ‘0.04 uM’, “2” = ‘0.12 uM’, “3” = ‘0.37 uM’, “4” = ‘1.11 uM’, “5” = ‘3.33 uM’, “6” = ‘10 uM’). We first applied PCA to the LINCS dataset with QC (post-QC) or without (pre-QC) using the top 50 components, representing 85.60% variance for the pre-QC profile and 86.78% variance for the post-QC profile. We then applied UMAP for dimensionality reduction using 20 nearest neighbors and a minimum distance of 0.1 to balance local and global structure preservation.

#### Supervised MOA classification

We trained multi-class neural network MLP classifiers to predict 439 MOA classes from well-level aggregated spherized profiles using sklearn.neural_network. We included the same samples in the training and testing splits as the original LINCS paper^17^. We trained four separate multiclass models, each performing the same MOA prediction task but with different input feature space: 1) pre-QC model, which uses the pre-QC aggregated spherized feature selected feature space; 2) post-QC model, which uses the post-QC aggregated spherized feature selected feature space; and 3-4) two shuffled baseline models generated from the pre-QC and post-QC datasets separately. We generated shuffled baseline models by independently permuting each feature column to destroy the feature-label structure. For all models, we performed a grid search over hidden-layer architectures (e.g., a single layer of 64 units, a single layer of 128 units, or two layers of 128 and 64 units) using 5-fold stratified cross-validation, with model performance scored by micro-averaged F1. We trained each model with a maximum solver iteration limit of 200 during the search. We kept all other hyperparameters at sklearn default values, observing minimal impact in performance. Finally, we refit the optimal architecture per model on the respective training set using all folds and with a maximum solver iteration of 500. We used a random state of 0. We evaluated final model performance on the held-out test set using per-class precision and recall and area under the precision recall curve (AUPR), with only 222 MOAs appearing in the held out test set from the original 439 trained MOA classes.

#### Ranking compounds with mean average precision

We generated new well-level aggregated spherized feature-selected profiles by keeping only cells that failed coSMicQC using the same Pycytominer pipeline. We calculated the compound and dose-level mean average precision (mAP) scores from the copairs^45^ software (using default parameters) for the three sets of bulk profiles; 1) all cells (pre-QC), only cells passing coSMicQC (post-QC), and only cells that failed coSMicQC. We used the aggregated profiles from DMSO-control wells as reference, specifically only comparing the DMSO-wells from the plates per compound+dose. The reference is built separately pre-QC and post-QC. For the failing cell only bulk profile, we used passing-cell DMSO as a fixed reference to isolate the effect of treatment-cell quality on mAP, since failing-QC DMSO wells are themselves expected to resemble failing-QC treatment wells rather than serving as an independent baseline as we applied coSMicQC to all wells. We assigned compound rankings based on mAP value, with compounds at mAP score of 1 will be assigned to rank 1. We computed compound rankings independently per dose. To identify compounds rescued by QC, we selected those that entered the top 20 ranking post-QC for a given dose but were not in the top 20 pre-QC. Of the rescued compounds+doses, we computed pairwise cosine similarities per replicate for pre and post-QC profiles.

#### Evaluating the impact of coSMicQC on lead compound nomination

For the compounds that were either rescued or left top-20 ranking, we computed pairwise cosine similarity of the replicates per compound+dose and compared to DMSO replicates from the same plates pre and post-QC. We calculated the average of the cosine similarity within replicates and compared to DMSO and computed the change in cosine similarity from pre-QC to post-QC. For all compounds, we calculated the mean fraction of single-cells that failed QC per compound+dose and took the number of cells that flagged QC per well over the total number of segmented cells and took the average of those fractions. We also calculated average replicate cosine similarity scores across replicate wells for each compound and doses for both bulk aggregate profiles (pre- and post-QC). We split compounds into groups based on if the mAP increased or decreased after QC was applied (only passing cells mAP scores - all cells mAP scores) to compare how similar replicates are between cells that failed QC or passed QC. We computed the t-statistic of the cosine similarity scores per dose between compound distributions that increased mAP score after coSMicQC was applied versus compounds that decreased mAP. As well, we compared mAP scores derived from failing-cells-only profiles against the original mAP scores derived from either all cells (no QC) or passing cells only (coSMicQC applied). Compound + doses with no change in mAP score (Δ = 0) were excluded from the visualizations.

## Acknowledgements

We would like to thank Vince Rubinetti for their work on logo generation for coSMicQC. We also thank Mike Lippincott, Cameron Mattson, Erik Serrano, Julia Curd, and Roshan Kern for their contributions in code review. We are grateful to our collaborators for their work collecting the image sets used in the development of coSMicQC. Research reported in this publication was supported by The Gilbert Family Foundation (923014 to G.P.W.), Alex’s Lemonade Stand Foundation ‘A’ Award and Tap Cancer Out (Grant # 23–28306 to G.P.W.), and American Heart Association Collaborative Sciences Award (24CSA1255857 to G.P.W.).

## Code availability

The coSMicQC software^36^ can be found on GitHub within the Cytomining organization. All code used to generate the figures can be found in the documentation (“docs”) folder of the repository.

