## Supplementary Figures for "Stellar quality control for single-cell image-based profiling with coSMicQC"

A

**Condition: Find over-segmented nuclei (e.g., clusters)**

```
nuclei_clustered_outliers = find_outliers(
    df=filtered_plate_df,
    metadata_columns=metadata_columns,
    feature_thresholds={
        "Nuclei_Intensity_MassDisplacement_CorrDNA": 0.05,
        "Nuclei_Intensity_IntegratedIntensity_CorrDNA": 1.5,
    },
)
```

**Create CytoDataFrame to visualize outliers**

```
nuclei_clustered_outliers_cdf = CytoDataFrame(
    data=pd.DataFrame(nuclei_clustered_outliers), # Convert back to pandas to apply transformations
    data_outline_context_dir="/path/to/outlines",
    segmentation_file_regex=outline_to_orig_mapping,
    display_options={
        "center_dot": False,
        "outline_color": (180, 30, 180),  # magenta
        "brightness": 20,
    },
)[
    [
        "Nuclei_Intensity_MassDisplacement_CorrDNA",
        "Nuclei_Intensity_IntegratedIntensity_CorrDNA",
        "Image_FileName_OrigDNA",
    ]
]
```

B

```
Number of outliers: 3630 (4.62%)
Outliers Range:
Nuclei_Intensity_MassDisplacement_CorrDNA Min: 0.5999707298906707
Nuclei_Intensity_MassDisplacement_CorrDNA Max: 14.840765847957671
Nuclei_Intensity_IntegratedIntensity_CorrDNA Min: 389.6632658392191
Nuclei_Intensity_IntegratedIntensity_CorrDNA Max: 1757.922207608819
(3630, 3)
```

C

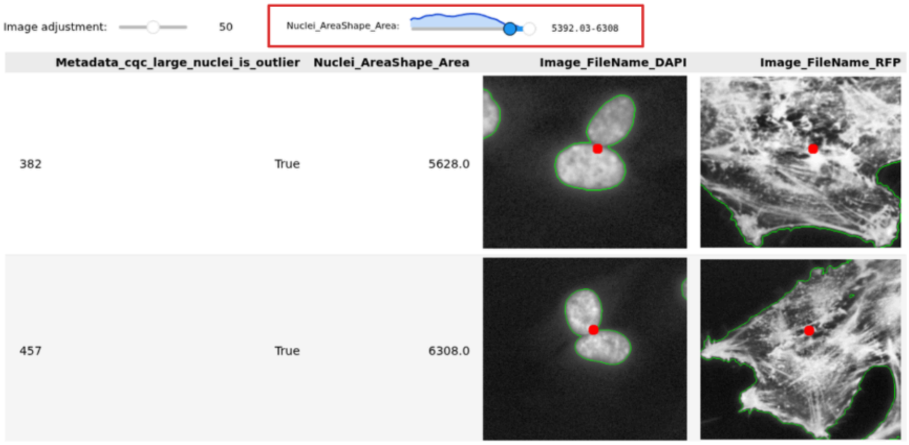

Supplemental Figure 1: Implementation of coSMicQC and CytoDataFrame for detecting outlier segmentations

(**A**) Example code to set one condition that detects over-segmented nuclei using coSMicQC (v1.1.0) and converting the output to a CytoDataFrame (v0.3.2). In this case, the condition uses two CellProfiler features: Nuclei_Intensity_MassDisplacement_CorrDNA and Nuclei_Intensity_IntegratedIntensity_CorrDNA, and sets thresholds of 0.05 and 1.5 standard deviations, respectively. CytoDataFrame has many display options to optimize visualizing the single-cell crops. (**B**) Example report output from coSMicQC with the summary statistics of technical outliers. (**C**) Example CytoDataFrame output in a Jupyter-based notebook, which visualizes each crop and corresponding mask outlines per row next to the respective feature values. In this example, a user would discard these undersegmentation of nuclei. Image brightness can be adjusted on the fly, and the dataframe can be filtered based on the distribution of the feature

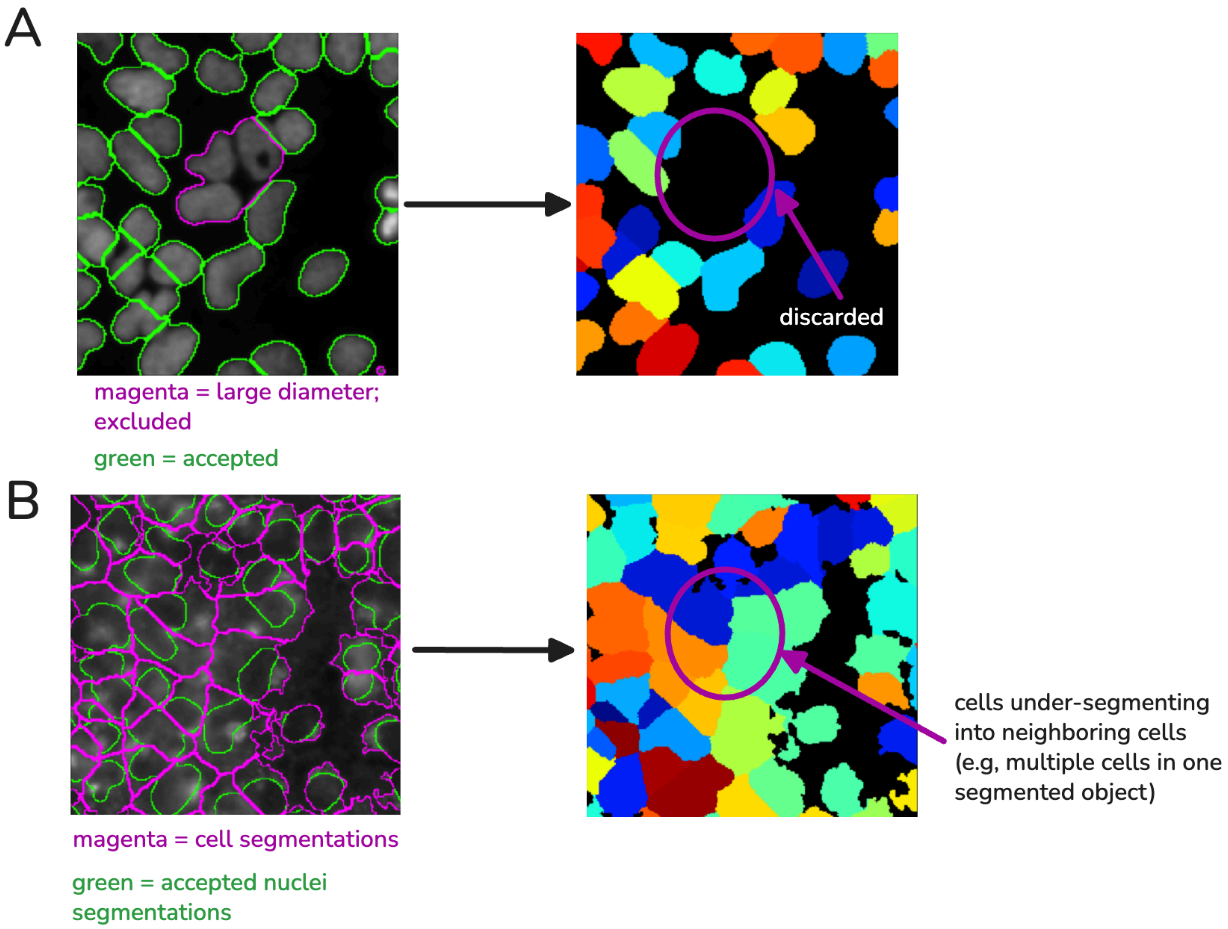

Supplemental Figure 2: Example under-segmented cells that results from pre-filtering of nuclei

(**A**) Example segmentation, which shows an under-segmented cluster of nuclei (magenta). Segmentation tools (e.g., CellProfiler) will filter this object because of a preset parameter that indicates an oversized diameter (e.g., based on parameters in IdentifyPrimaryObjects). Because popular algorithms like Otsu thresholding seed nuclei masks to find individual cell objects, these algorithms tend to undersegment the cells of neighboring discarded nuclei. (**B**) The region that would have contained the discarded nuclei is inappropriately segmented as regions of neighboring cells. Including these cells in single-cell analyses, will contaminate findings.

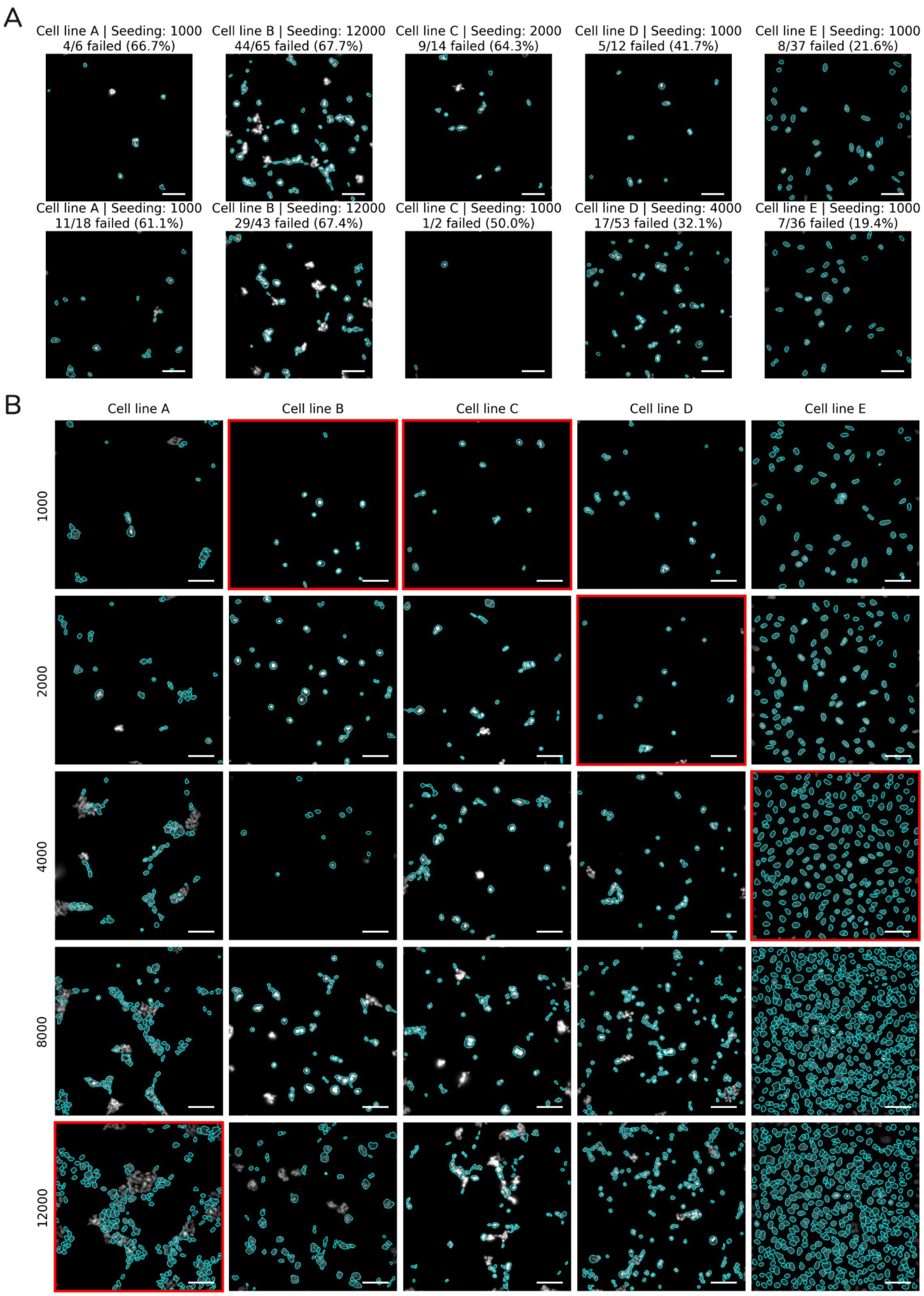

Supplemental Figure 3: coSMicQC detects bad nuclear segmentations across cell lines to help optimize cell line seeding density

(**A**) Example FOVs with the highest fraction of cells failing coSMicQC per cell line shows coSMicQC can detect instances of low quality segmentations in non-optimal conditions. (**B**) Example FOVs from each cell line (x-axis) and seeding density (y-axis), with red boxes over the seeding density with the lowest coSMicQC failure rate. Nuclei segmentation outlines are in cyan. The scale bar is 100 uM. The original cell lines' names in order from A-E are as follows: CHLA-10, CHLA-113, CHLA-218, CHLA-25, and U2-OS.

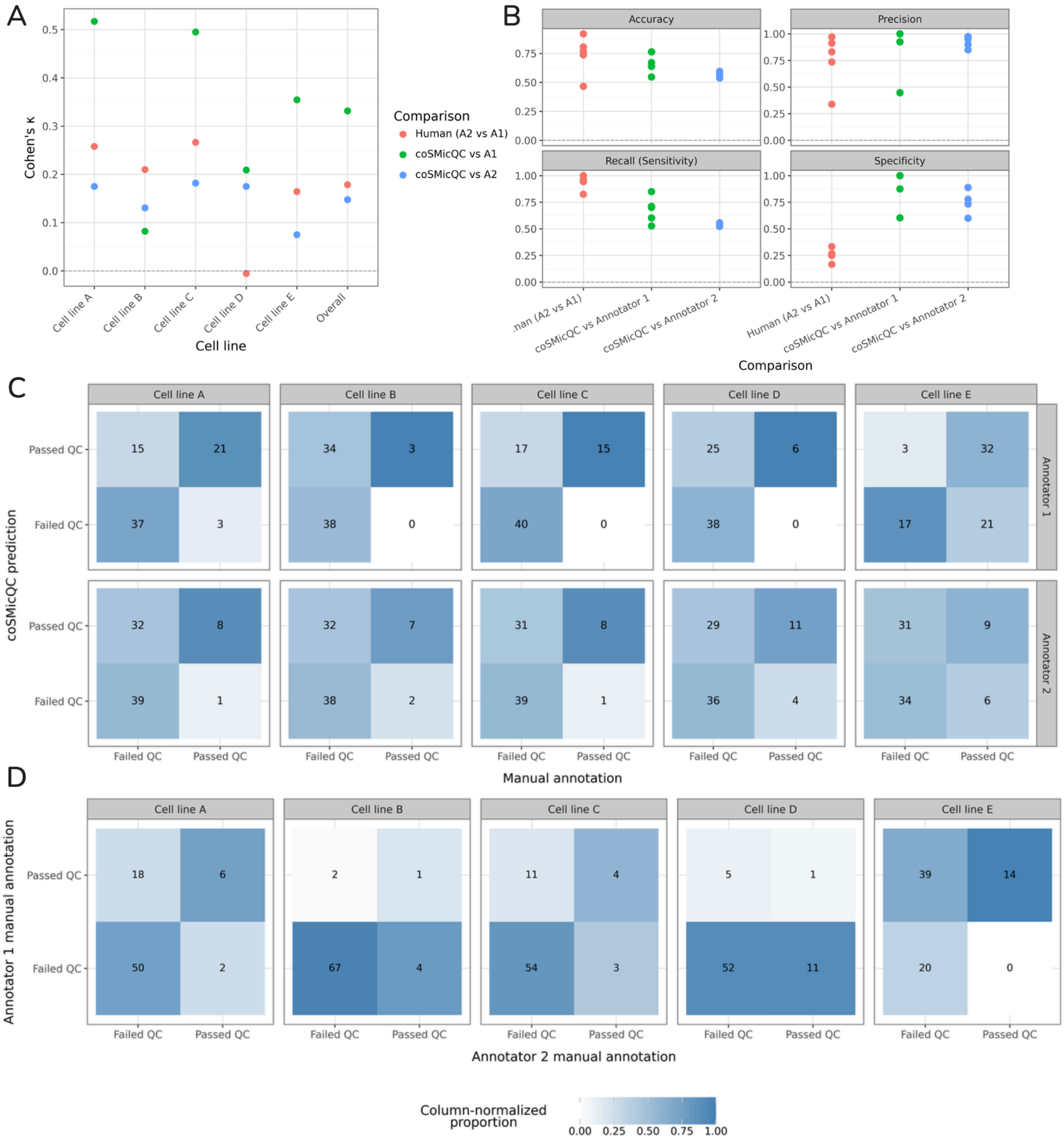

### Supplemental Figure 4: coSMicQC demonstrates high specificity but reduced sensitivity in detecting poor-quality segmentations compared to manual annotation

(**A**) Cohen's kappa agreement scores between two human annotators and coSMicQC across cell lines and overall. CoSMicQC consistently falls either within the agreement range for human annotators or above. (**B**) Performance metrics between coSMicQC and the human annotators compared to the human annotators only. Each point represents a cell line. The named annotating method (i.e., A1, A2, or coSMicQC) listed second is considered ground truth for the comparison. Confusion matrices of (**C**) coSMicQC and manual annotations across two annotators and five cell lines and (**D**) each of the two annotators annotations across cell lines. The original cell lines' names in order from A-E are as follows: CHLA-10, CHLA-113, CHLA-218, CHLA-25, and U2-OS.

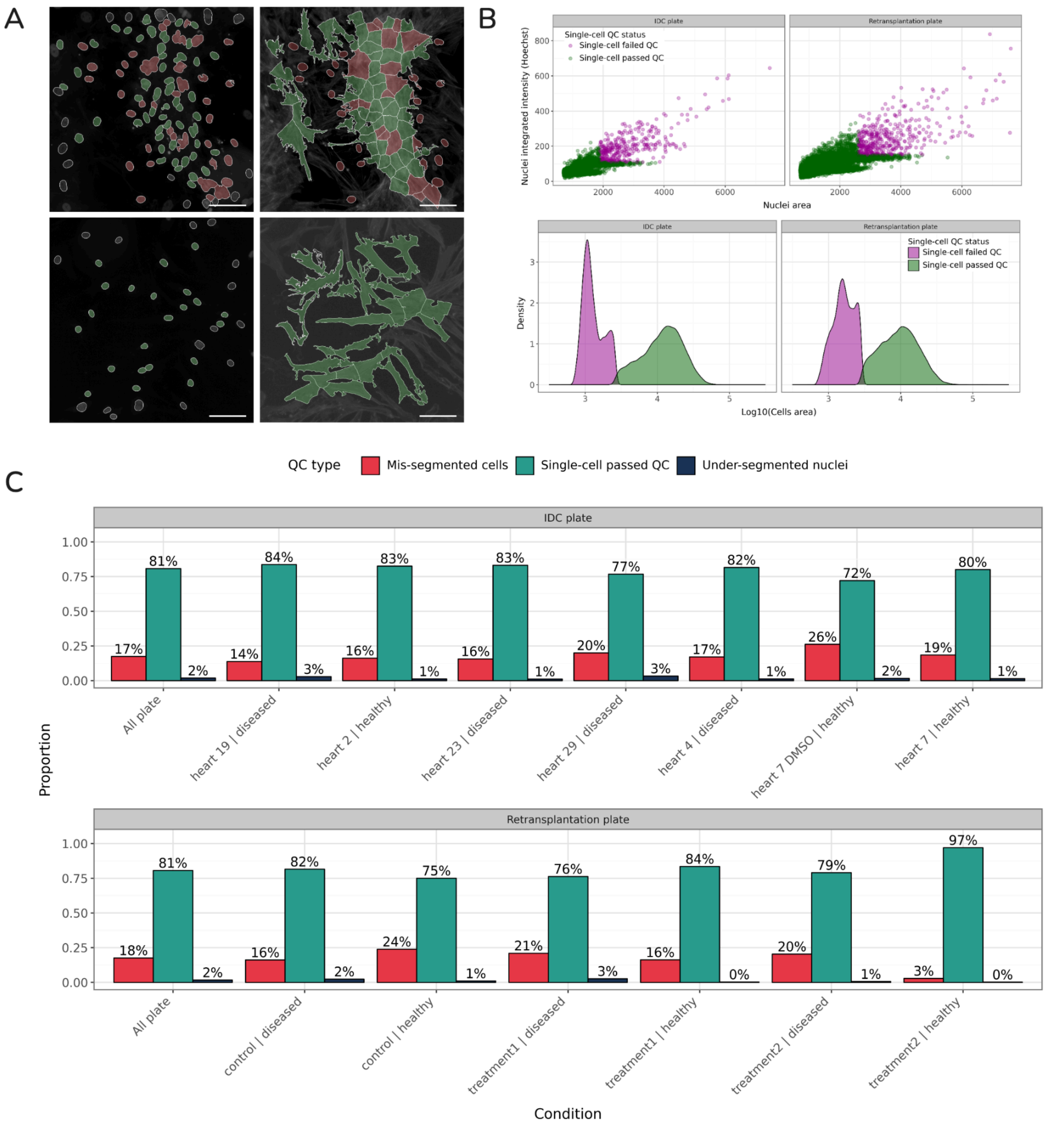

Supplemental Figure 5: Quantifying outlier removal in plates from the cardiac fibrosis dataset

(**A**) Cells that fail quality control conditions (red) can be seen in the same field of view (FOV) as cells that are properly segmented (green). Segmentations in grey represent cells that are touching an edge and are discarded from the profile. The scale bar is 200 μM. There are many cases where nuclei are under-segmented or cells are mis-segmented due to intensity issues associated with higher confluency and lack of contact inhibition. (**B**) CoSMicQC detects abnormally small cells using cell area to identify debris or other segmentation artifacts arising from high confluence. We used slightly different thresholds between plates (Δ = 0.1). (**C**) The rate of detected outliers across conditions is consistent across treatments and cell types, with an average of over 80% of cells passing QC.

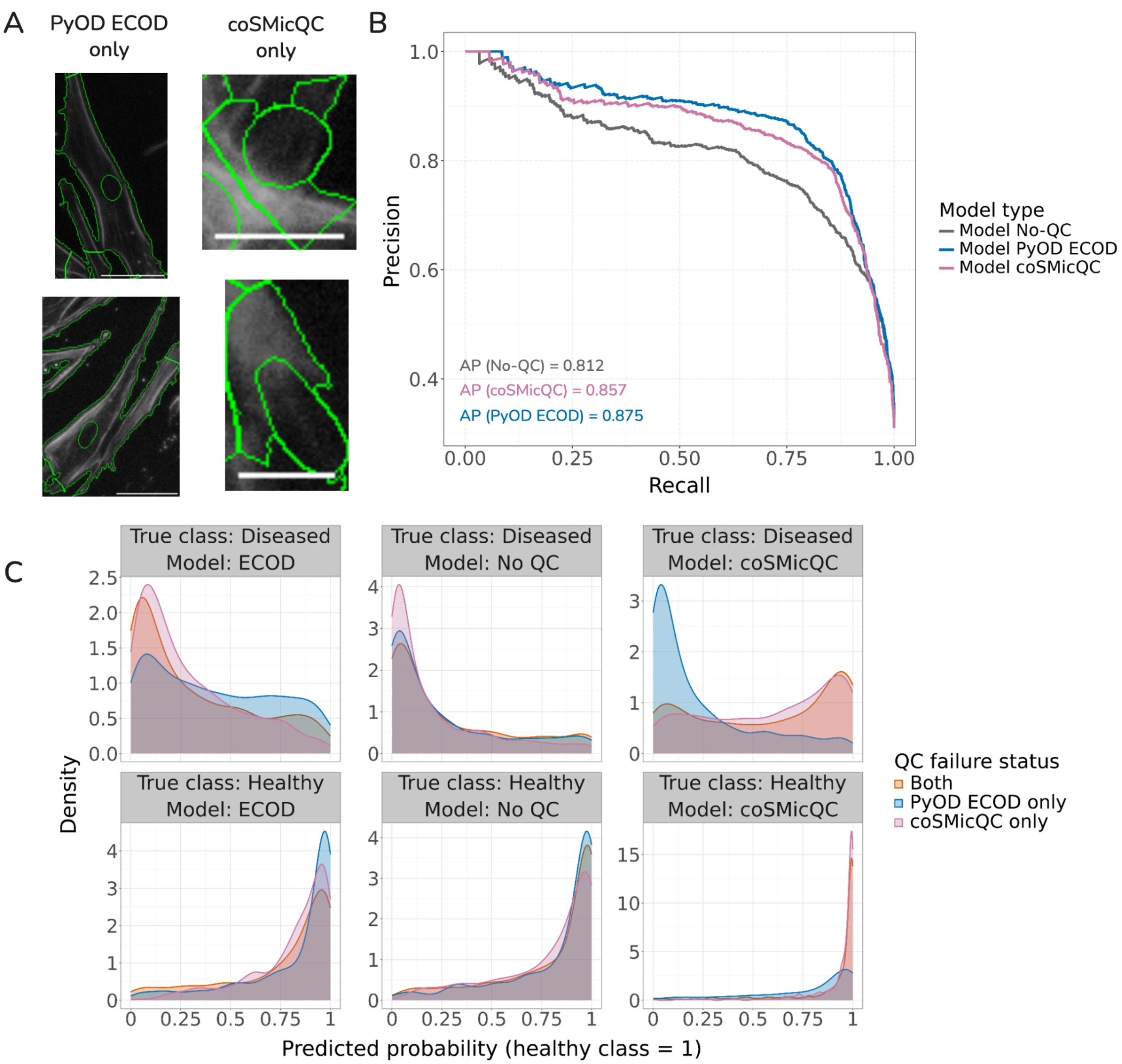

Supplemental Figure 6: CoSMicQC removes segmentation artifacts and increases model performance similarly to PyOD ECOD

(**A**) Example failing cells from each QC method from the Treatment 2 healthy heart from the retransplantation plate. The scale bar is 100 μM. Cells failing PyOD ECOD do not always reflect the same segmentation errors that coSMicQC flags as failing cells. (**B**) Precision-recall curves of each model applied to a holdout set of cells from the IDC plate (DMSO-treated healthy heart and one failing heart) demonstrate the ECOD model performs slightly better than coSMicQC, but both perform better than if no QC is applied at all. (**C**) Predicted probability distributions on cells failing the specified QC method. The facet labels indicate the model type (which data we trained the model using) and the class of the cells that failed QC.

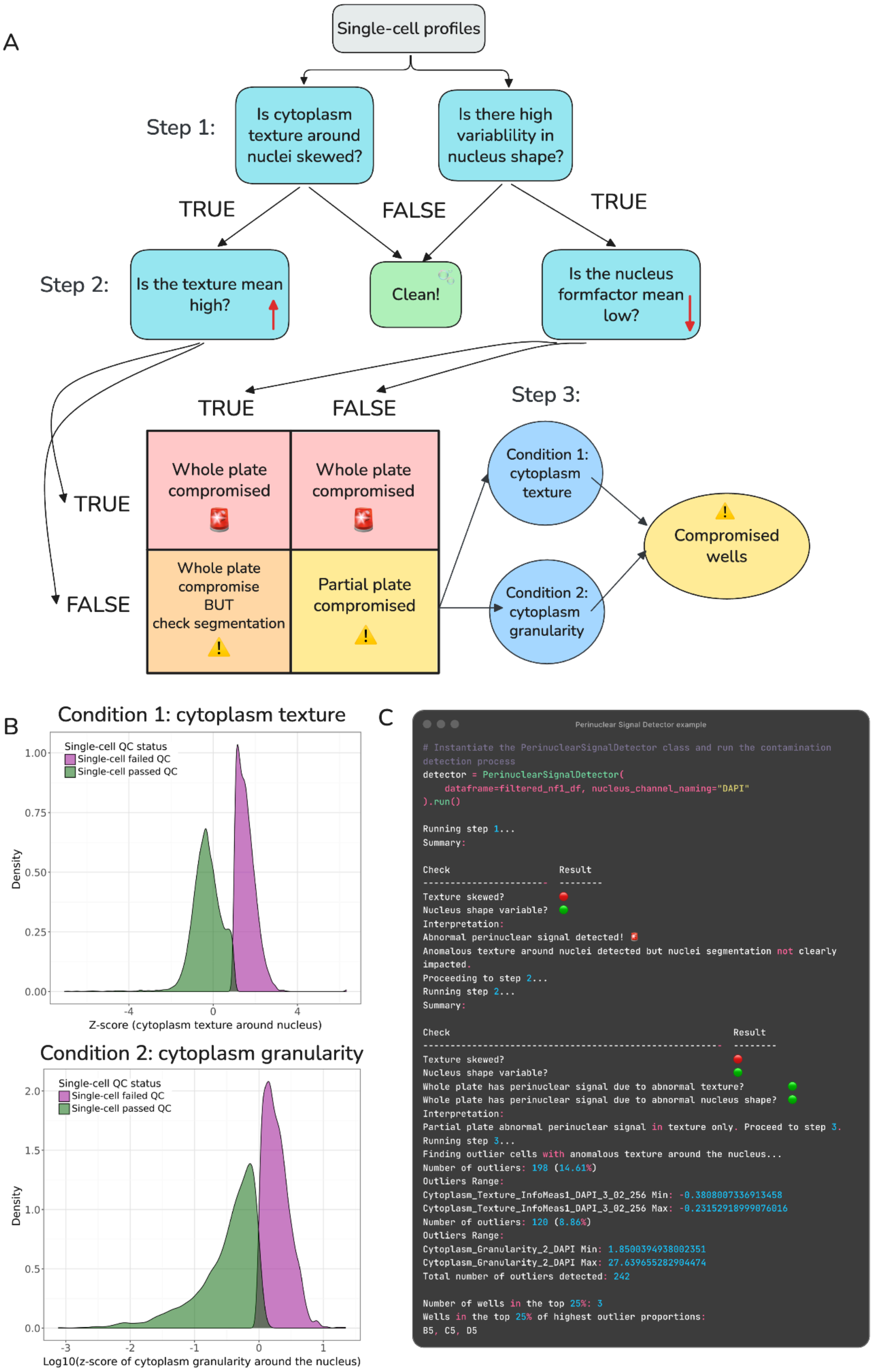

### Supplemental Figure 7: The coSMicQC perinuclear signal detector

(**A**) A decision tree detailing the three-step perinuclear signal detector in coSMicQC. In the first step, the detector answers two questions. The answers to these questions will determine if the plate is clean or if we move on to step two (see main text for details). In the second step, the detector will answer two additional questions, and the answers lead to the 2x2 grid. It is unlikely that the answer to the first and second questions is False and True, respectively, but we have found that this combination likely indicates poor segmentation rather than a whole plate being compromised (orange square). In the third step, the detector uses the standard coSMicQC approach, setting two conditions to identify compromised single cells (partial contamination; some wells and not the whole plate). (**B**) We present real-world examples of the distributions for the conditions in Step 3. The (top) cytoplasm texture condition and (bottom) cytoplasm granularity condition clearly distinguish contaminated cells. (**C**) Example API usage for perinuclear signal detector with output for partial plate detection.

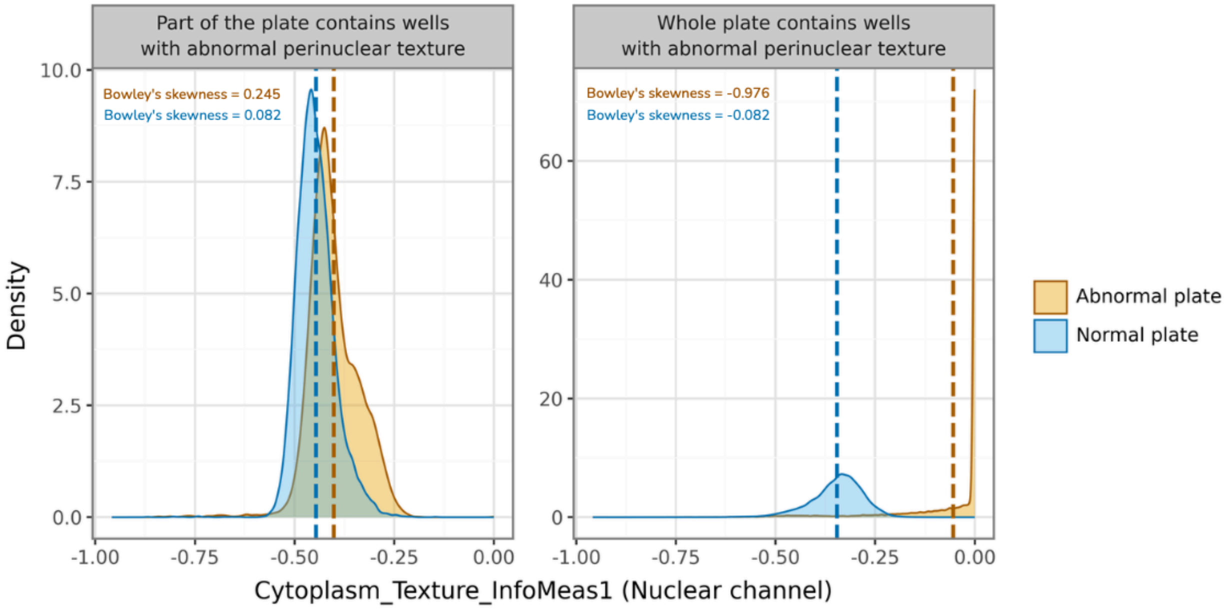

Supplemental Figure 8: Abnormal perinuclear signal skews cytoplasm texture in the nuclear channel

Plates with abnormal perinuclear texture distributions (orange) show left skewness compared to plates without the abnormal texture (blue), which can be detected by quantitative measurements like Bowley’s skewness. In some cases, (left) only certain wells within a plate contain abnormal signal; (right) in others, the entire plate is affected, resulting in extreme skewness. The dotted lines represent the mean of the distributions.

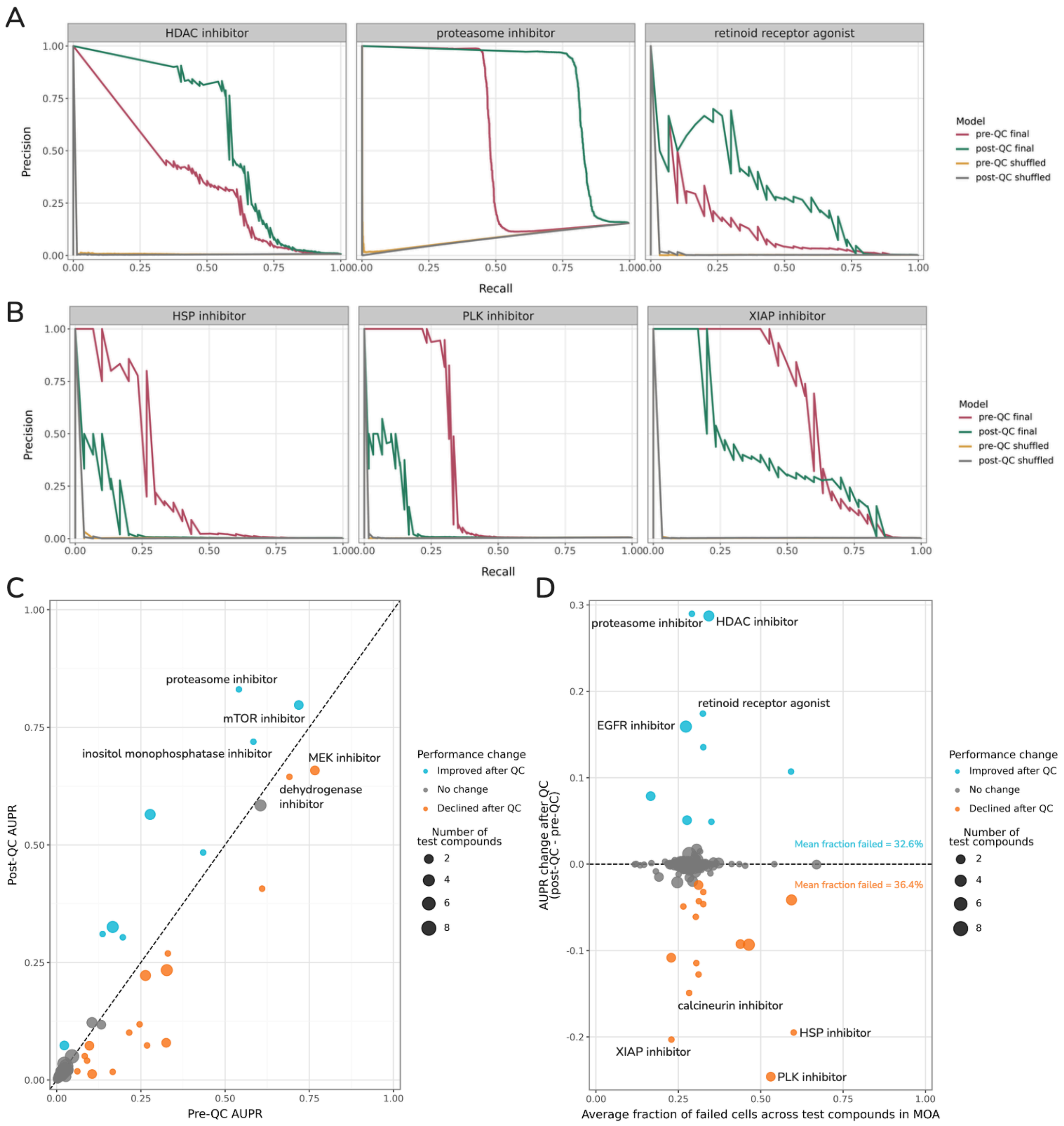

### Supplemental Figure 9: CoSMicQC marginally impacts MOA classification

Test-set precision-recall curves for (**A**) the top three MoAs with the highest increase in performance after quality control, and (**B**) the bottom three performing MOAs after coSMicQC. (**C**) Comparing test-set AUPR scores from the pre- and post-QC model for each MOA. (**D**) coSMicQC flags cells at a similar rate across compounds with no meaningful relationship to MOA prediction performance.

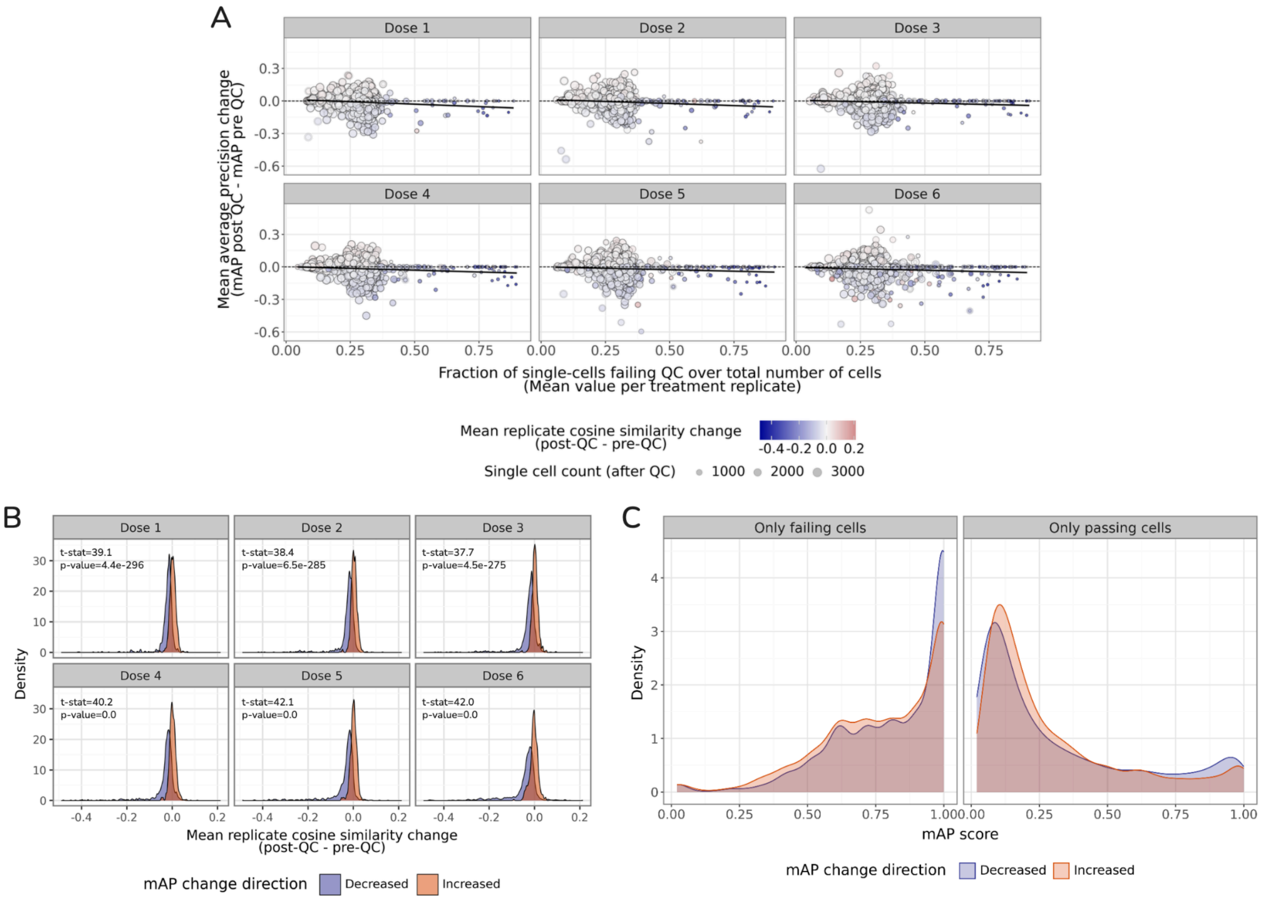

Supplemental Figure 10: The nuanced relationship between mAP and coSMicQC reveals that technical outliers are similar to each other and, when removed from the profiles, sometimes results in treatments with lower performance after QC

(**A**) Mean average precision (mAP) change (post-QC minus pre-QC) compared to the fraction of single cells failing QC across compounds and doses. Compounds with high amounts of cells removed have generally lower mAP scores after QC. (**B**) Compounds with increased mAP after QC either had no change or slight increase in cosine similarity between replicates. Compounds with decreased mAP showed a drop in cosine similarity between replicates. We calculated replicate correlation by applying cosine similarity to pre- and post-QC profiles. (**C**) Aggregated profiles generated from compound-treated cells passing QC had substantially lower mAP than profiles generated from failing cells, indicating that cells failing QC are similar to each other and, in some compounds, QC is likely to reduce mAP.

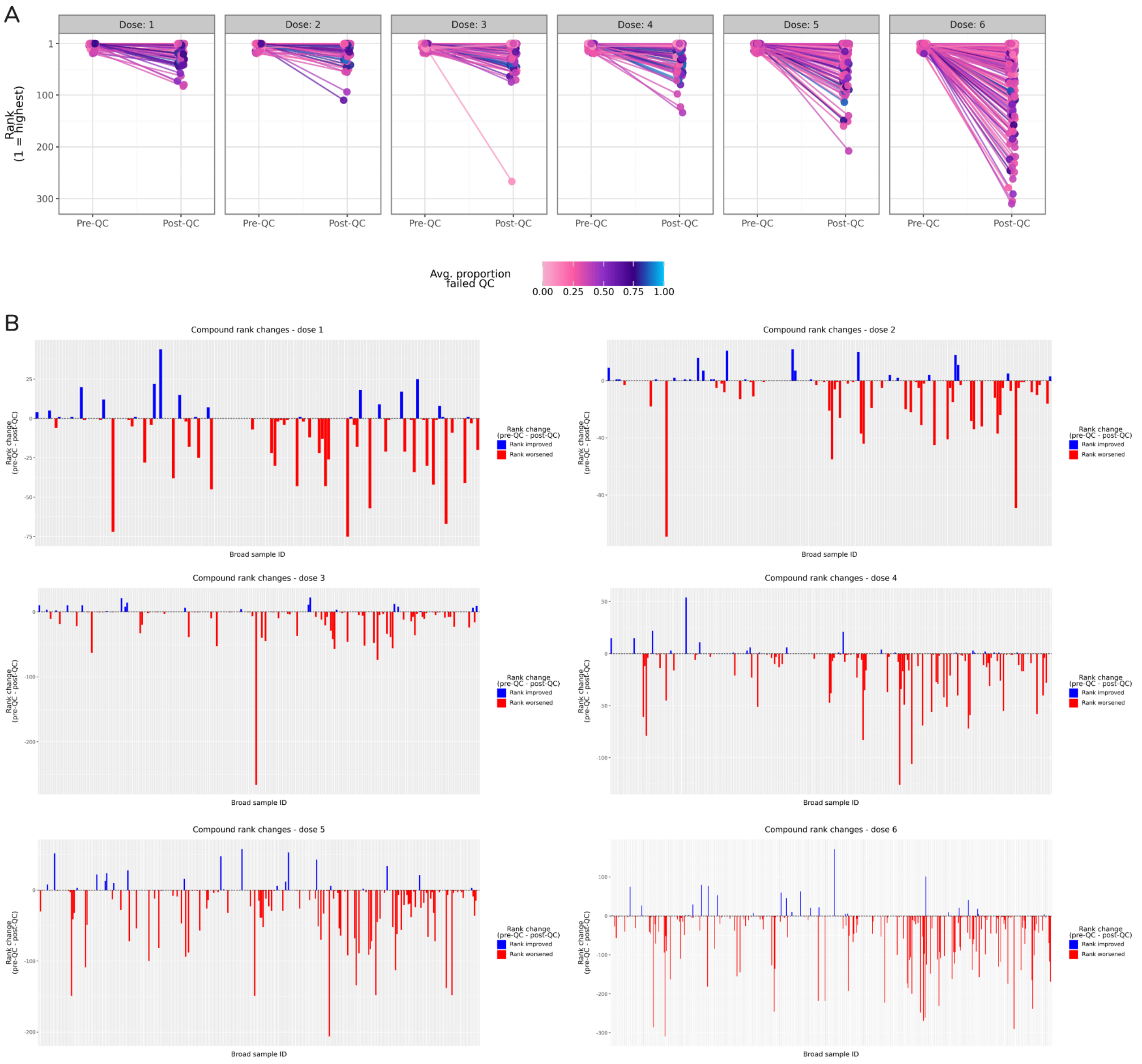

### Supplemental Figure 11: Rank changes for lead compounds in LINCS

(**A**) Quality control impacts lead compound ranking across doses. Each line is a compound and is found in the top-20 ranking pre-QC per dose. (**B**) All six doses showed changes in compounds within the top 20 rankings after quality control. Blue indicates an improvement in rank, while red indicates a lowered rank number (i.e., the compound moved down and had a lower mAP score).

Supplemental Figure 12: Rescued treatments show minimal change in cosine similarity after quality control

(**A**) Pairwise cosine similarity heatmap of the 50 rescued treatments pre-QC. (**B**) Pairwise cosine similarity heatmap of the 50 rescued treatments post-QC. Both heatmaps are ordered by compound, MOA, and dose. (**C**) Distributions of pairwise cosine similarities demonstrates no difference between pre- and post-QC.

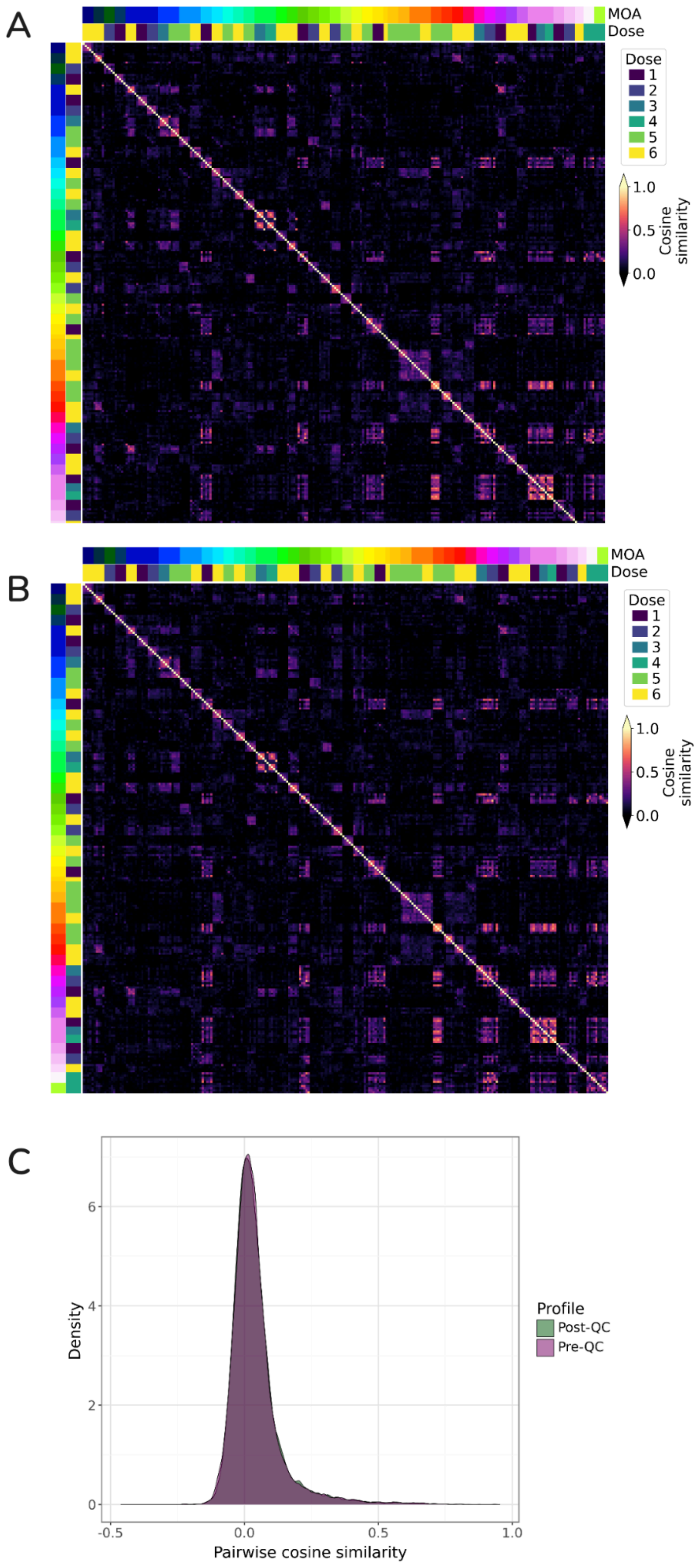
